# A comprehensive proteomic and bioinformatics analysis of human spinal cord injury plasma identifies proteins associated with the complement cascade and liver function as potential prognostic indicators of neurological outcome

**DOI:** 10.1101/2022.07.12.499696

**Authors:** Gabriel Mateus Bernardo Harrington, Paul Cool, Charlotte Hulme, Aheed Osman, Joy Roy Chowdhury, Naveen Kumar, Srinivasa Budithi, Karina Wright

**Affiliations:** Keele University, Staffordshire, United Kingdom, ST5 5BG; Cardiff University, Cardiff, United Kingdom, CF24 4HQ; Robert Jones and Agnes Hunt Orthopaedic Hospital NHS Foundation Trust, Oswestry, United Kingdom, SY10 7AG

**Keywords:** Spinal cord injury, Biomarker, Proteomics, Complement cascade

## Abstract

1.

**Introduction:** Spinal Cord Injury (SCI) is a major cause of disability, with complications post-injury often leading to life-long health issues with need of extensive treatment. Neurological outcome post-SCI can be variable and difficult to predict, particularly in incomplete injured patients. The identification of specific SCI biomarkers in blood, may be able to improve prognostics in the field. This study has utilised proteomic and bioinformatics methodologies to investigate differentially expressed proteins in plasma samples across human SCI cohorts with the aim of identifying prognostic biomarkers and biological pathway alterations that relate to neurological outcome.

**Methods and Materials:** Blood samples were taken, following informed consent, from ASIA impairment scale (AIS) grade C “Improvers” (those who experienced an AIS grade improvement) and “Non-Improvers” (No AIS change), and AIS grade A and D at <2 weeks (“Acute”) and approx. 3 months (“Sub-acute”) post-injury. The total protein concentration from each sample was extracted, with pooled samples being labelled and non-pooled samples treated with ProteoMiner™ beads. Samples were then analysed using two 4-plex isobaric tag for relative and absolute quantification (iTRAQ) analyses and a label-free experiment for comparison, before quantifying with mass spectrometry. Data are available via ProteomeXchange with identifiers PXD035025 and PXD035072 for the iTRAQ and label-free experiments respectively.

Proteomic datasets were analysed using OpenMS (version 2.6.0). R (version 4.1.4) and in particular, the R packages MSstats (version 4.0.1) and pathview (version 1.32.0) were used for downstream analysis. Proteins of interest identified from this analysis were further validated by enzyme-linked immunosorbent assay (ELISA).

**Results:** The data demonstrated proteomic differences between the cohorts, with the results from the iTRAQ approach supporting those of the label-free analysis. A total of 79 and 87 differentially abundant proteins across AIS and longitudinal groups were identified from the iTRAQ and label-free analyses, respectively. Alpha-2-macroglobulin (A2M), retinol binding protein 4 (RBP4), serum amyloid A1 (SAA1), Peroxiredoxin 2, Apolipoprotein A1 (ApoA1) and several immunoglobulins were identified as biologically relevant and differentially abundant, with potential as individual prognostic biomarkers of neurological outcome. Bioinformatics analyses revealed that the majority of differentially abundant proteins were components of the complement cascade and most interacted directly with the liver.

**Conclusions:** Many of the proteins of interest identified using proteomics were detected only in a single group and therefore have potential as a binary (present or absent) biomarkers, RBP4 and PRX-2 in particular. Additional investigations into the chronology of these proteins, and their levels in other tissues (cerebrospinal fluid in particular) are needed to better understand the underlying pathophysiology, including any potentially modifiable targets. Pathway analysis highlighted the complement cascade as being significant across groups of differential functional recovery.

## 2. Introduction

Spinal cord injury (SCI) is the transient or permanent loss of normal spinal sensory, motor or autonomic function, and is a major cause of disability. Globally, SCI affects around 500,000 people each year and is most commonly the result of road traffic accidents or falls. ^1^ Patients typically require extensive medical, rehabilitative and social care at high financial cost to healthcare providers. The lifetime cost of care in the UK is estimated to be £1.12 million (mean value) per SCI, with the total cost of SCI in the UK to the NHS being £1.43 billion in 2016. ^2^ Individuals with SCI show markedly higher rates of mental illness relative to the general population. ^3^ Complications arising post-SCI can be long-lasting and often include pain, spasticity and cardiovascular disease, where the systemic inflammatory response that follows SCI can frequently result in organ complications, particularly in the liver and kidneys. ^4,5,6^

The recovery of neurological function post-SCI is highly variable, requiring any clinical trials to have an impractically large sample size to prove efficacy, hence the translation of novel efficacious therapies is challenging and expensive. ^7^ Being able to more accurately predict patient outcomes would aid clinical decisions and facilitate future clinical trials. Therefore, novel biomarkers that allow for stratification of injury severity and capacity for neurological recovery would be of high value to the field.

Biomarkers studies in SCI often investigate protein changes in cerebral spinal fluid (CSF) as the closer proximity of this medium is thought to be more reflective of the parenchymal injury. ^8,9^ Whilst this makes CSF potentially more informative for elucidating the pathology of SCI, the repeated use of CSF for routine analysis presents challenges in clinical care due to the risk and expense associated with the invasiveness of the collection procedure. In contrast, systemic biomarkers measurable in the blood represent a source of information that can be accessed and interpreted both a lower cost and risk. Studies of traumatic brain injury have demonstrated that protein markers identified in CSF are also detectable in both plasma and serum. ^10^ More recently, circulating white blood cell populations have also been identified as potential SCI injury biomarkers, with a 2021 study showing that elevated levels of neutrophils were associated with no AIS grade conversion, while conversely an increase in lymphocytes during the first week post-SCI were associated with an AIS grade improvement.^11^

A number of individual proteins have been shown to be altered in the bloods post-SCI, including multiple interleukins (IL), tumour necrosis factor alpha (TNF-*α*) and C-reactive protein (CRP). ^12,13,14^

Further, changes in inflammatory marker levels detected in acute SCI patients were found to be mirrored in donor-matched blood and CSF, albeit at lower absolute concentrations systemically.^15^Previously, we have shown that routinely collected blood measures associated with liver function and inflammation added predictive value to AIS motor and sensor outcomes at discharge and 12-months post-injury. ^16,17^ The current study uses an unbiased shotgun proteomic approach to investigate differentially expressed proteins in SCI patients, coupled with bioinformatics pathway and network analyses.

## 3. Methods and Materials

### 3.1. Patients

Blood samples were taken from SCI patients who had provided informed consent and in accordance to ethical provided by the National Research Ethics Service (NRES) Committee North West Liverpool East (11/NW/0876). “Improvers” were defined as individuals who experienced an AIS grade improvement from admission to a year post-injury, whereas “non-improvers” were defined as patients who saw no change in AIS grade in the same period (Table 1).

**Table 1:**
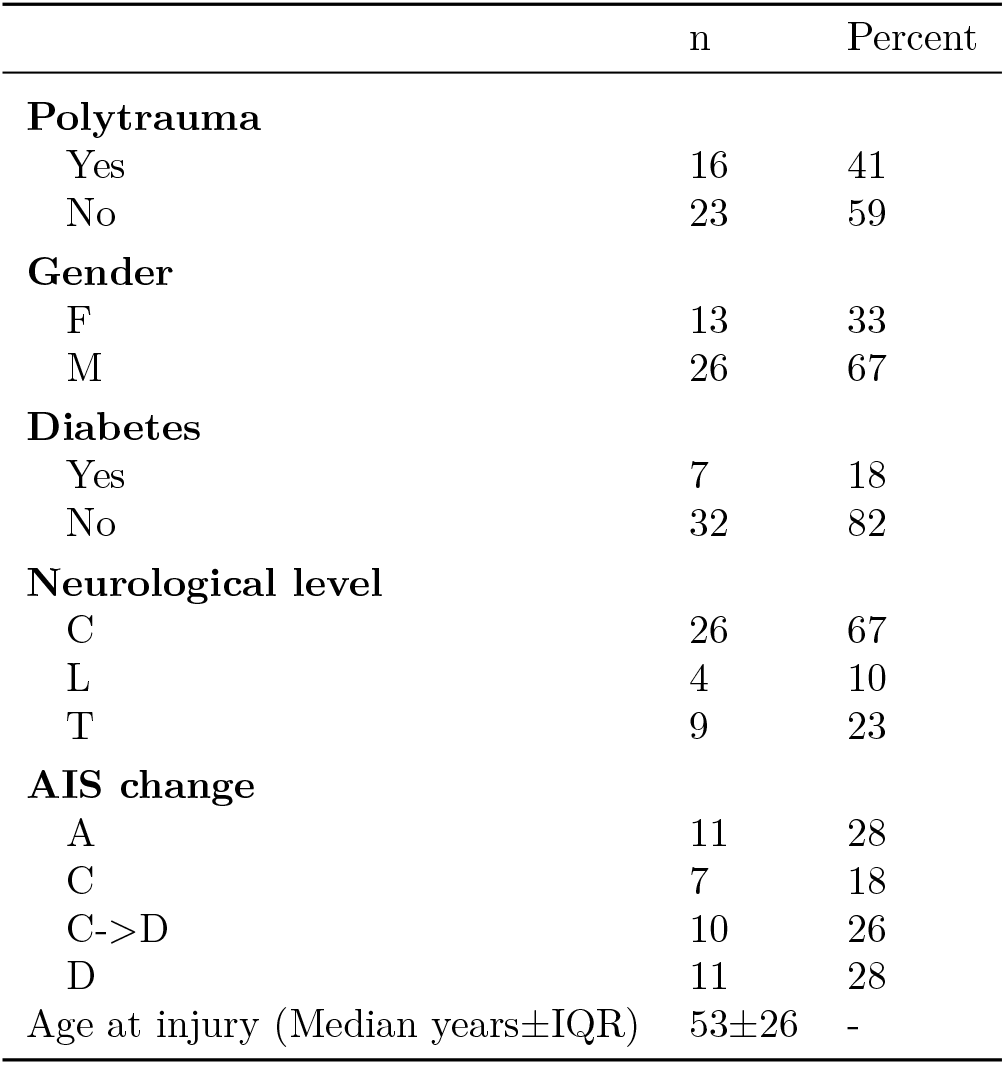
Patient demographics. ± denotes interquartile range

### 3.2. Plasma collection and storage

Plasma samples were collected within 2 weeks of injury (acute) and at approximately 3 months post-injury (subacute). Upon collection in EDTA (ethylenediaminetetraacetic acid) coated tubes samples were centrifuged at 600g for 15 minutes, to pellet erythrocytes and the resultant plasma fraction was aspirated and divided into aliquots for long-term storage in -80°C briefly and liquid nitrogen in the longer term.

### 3.3. Sample preparation and analysis using iTRAQ proteomics

Thawed plasma samples (2*µl*) each were diluted with distilled water (98*µl*). Total protein was quantified using a Pierce™ 660*nm* Protein Assay (Thermo Fisher Scientific, Hemel Hempstead, UK) ^18^.

A total of 100*mg* of plasma protein was taken from each sample and pooled equally to form a patient test group. For example, the AIS C improver group was pooled from 10 separate patient samples, 10mg of protein per patient.

The pooled plasma samples were precipitated by incubation of the sample in six times the volume of chilled acetone for 1 hour at -20°C. The samples were then centrifuged at 6,000G for 10 minutes at 4°C, and re-suspended in 200*µl* of triethylammonium bicarbonate buffer. Sequencing Grade Modified Trypsin (10*µg/*85*µg* of protein; Promega, Madison, WI, USA) was then added to the samples for overnight digestion at 37°C. Peptides underwent reduction and alkylation (according to the manufacturer’s instructions; Applied Biosystems, Bleiswijk, The Netherlands). Tryptic digests were labelled with iTRAQ tags (again according to the manufacturer’s instructions for the iTRAQ kit), before being pooled into test groups and dried in a vacuum centrifuge. Two individual iTRAQ experiments were set up, the first to assess acute and sub-acute improvers or non-improvers and the second to assess acute improvers and non-improvers to AIS grade A and D patients. The following tags were used for each group of patient samples 114 tag -acute improvers, 115 tag - sub-acute improvers, 116 tag - acute non-improvers and 117 tag - sub-acute non-improvers for run 1 and 114 tag - acute improvers, 115 tag - acute non-improvers, 116 tag - AIS grade A and 117 tag - AIS grade D for run 2.

### 3.3.1. iTRAQ mass spectrometry analysis

The samples were analysed at the BSRC St. Andrews University Mass Spectrometry and Proteomics Facility. A total of 12 SCX fractions were analysed by nano-electrospray ionisation-liquid chromatography/tandem mass spectrometry (LC-MS/MS) using a TripleTOF 5600 tandem mass spectrometer (AB Sciex, Framingham, MA, USA) as described previously. ^19^ Each fraction (10*µl*) was then analysed by nanoflow LC-ESI-MSMS, as described previously.

The mass spectrometry proteomics data have been deposited to the ProteomeXchange Consortium via the PRIDE partner repository with the dataset identifier PXD035025 and 10.6019/PXD035025. ^20^

### 3.4. Sample preparation and analysis using label-free proteomics

No sample pooling was used, and so each of the 73 samples were maintained separately throughout protein equalisation, mass spectrometry, and label-free quantification steps. Thus, protein abundance was quantified for each sample, whereupon mean protein abundance across experimental groups was calculated to assess protein changes.

To reduce the dynamic range of proteins, ProteoMiner™ beads (BioRad, Hemel Hempstead, UK) were used. ^21^ Total protein was quantitated with a Pierce™ 660*nm* Protein Assay (Thermo Fisher Scientific, Hemel Hempstead, UK), whereupon 5 mg of total protein was applied to ProteoMiner™ beads, and processed as described previously. ^18,22^

#### 3.4.1. Label free mass spectrometry analysis

Tryptic peptides were subjected to LC-MC/MC via a 2-h gradient on a NanoAcquity™ ultraperformance LC (Waters, Manchester, UK) connected to a Q-Exactive Quadrupole-Orbitrap instrument (Thermo-Fisher Scientific Hemel Hempstead, UK).

The Q-Exactive was operated in a data dependent positive electrospray ionisation mode, automatically switching between full scan MS and MS/MS acquisition. Survey full scan MS spectra (*m/z* 300–2000) were acquired in the Orbitrap with 70,000 resolution (*m/z* 200) following accumulation of ions to 1 *×*10^6^ target value based on the predictive automatic gain control values from the previous full scan. Dynamic exclusion was set to 20s, the 10 most intense multiply charged ions (*z ≥*2) were sequentially isolated and fragmented in the octopole collision cell by higher energy collisional dissociation (HCD), with a fixed injection time of 100ms and 35,000 resolution. The following mass spectrometric conditions were used: spray voltage, 1.9kV, no sheath or axillary gas flow; normalised HCD collision energy 30%; heated capillary temperature, 250°C. MS/MS ion selection threshold was set to 1*×* 10^4^ count and 2Da isolation width was set.The mass spectrometry proteomics data have been deposited to the ProteomeXchange Consortium via the PRIDE partner repository with the dataset identifier PXD035072 and 10.6019/PXD035072. ^20^

### 3.5. iTRAQ OpenMS analysis

TripleTOF 5600 tandem mass spectrometer output files produced in the ABSciex proprietary .`wiff` file format were converted to an open file format, .`mzML` for analysis with OpenMS (version 2.6.0). The docker image of ProteoWizard version 3.0.20287 was used for conversion, and peak picking was applied on conversion ^23^. OpenMS version 2.6.0 was used for further analysis. ^24^ Unless otherwise stated, default arguments were used. The 12 fraction files were merged and sorted by retention time. A decoy database was generated with .`DecoyDatabase` and the .`-enzyme` flag set to `Trypsin`, the human reference proteome was taken from Uniprot (Proteome ID: UP000005640, downloaded: 2020-10-01), as was the .`fasta` for porcine trypsin (Entry: P00761, downloaded: 2020-10-01). ^25^

The `MSFGPlusAdapter` was used to run the search. For the `-fixed_modifications` “Methylthio (C)” and “iTRAQ4plex (N-term)” were passed due to the alkylating agent used in sample preparation and to account for the N-terminus modifications made by iTRAQ tags. “Oxidation (M)” was passed to `-variable_modifications` to reflect the likely occurrence of methionine oxidation. To reflect the instrument the following flags were also set: `-precursor_mass_tolerance 20 -enzyme Trypsin/P -protocol iTRAQ -instrument high_res`.

To annotate the search results `PeptideIndexer` and `PSMFeatureExtractor` were used. For peptide level score estimation and filtering `PercolatorAdapter` was used with the following arguments: `-score_type q-value -enzyme trypsinp. IDFilter` was used to filter to a peptide score of 0.05 with `-score:pep 0.05`

`IsobaricAnalyzer with -type itraq4plex` was used with the merged .`mzML` files to assign protein-peptide identifications to features or consensus features with `IDMApper`. The files for each run output by `IDMapper` were then merged with `FileMerger`. Bayesian score estimation and protein inference was performed with `Epifany` and the following flags: `-greedy_group_resolution remove_proteins_wo_evidence -algorithm:keep_best_PSM_only false` Decoys were removed and 0.05 FDR filtering was done via IDFilter with -score:protgroup 0.05 -remove_decoys. Finally, `IDConflictResolver` was used to resolve ambiguous annotations of features with peptide identifications, before quantification with `ProteinQuantifier`.

### 3.6. Label free OpenMS analysis

For quantification, the raw spectra files were analysed via OpenMS (version 2.6.0) command line tools, with the workflow from the prior section (Section 3.5) adapted to suit a label-free analysis. The files were first converted from the proprietary .Raw format to the open .mzML standard with the `FileConverter` tool via the open-source `ThermoRawFileParser`. ^24,26^ Unless otherwise stated, default arguments were used throughout.

The decoy database generated in the prior section (iTRAQ OpenMS analysis) was also re-used. The `CometAdapter` was used to run the search. ^27^ Fixed modifications were set to “Carbamidomethyl (C)” and “Oxidation (M)” was set as a variable modification. To reflect the instrument the following flags were also set: `-precursor_mass_tolerance 20 -isotope_error 0/1`.

To annotate the identified peptides with proteins the `PeptideIndexer` tool was used. `PeptideIndexer` and `PSMFeatureExtractor` were used for annotation. For peptide level score estimation and filtering `PercolatorAdapter` was used with the following flags: `-score_type q-value -enzyme trypsin. IDFilter` was used to filter to a pep-tide score of 0.01 with `-score:pep 0.01` followed by `IDScoreSwitcher` with the fol-lowing flags: `-new_score “MS:1001493” -new_score_orientation lower_better -new_score_type “pep” -old_score “q-value”`. `The ProteomicsLFQ` was used for subsequent processing with the flags: `-proteinFDR 0.05 -targeted_only true`. The -out_msstats flag was also used to produce quantitative data for downstream statistical analysis with the R package `MSstats`. ^28^

### 3.7. Network and pathway analysis

The Bioconductor package ReactomePA, which employs the open-source, open access, manually curated and peer-reviewed pathway database Reactome was used for network analysis. ^29,30^

### 3.8. Enzyme-linked immunosorbent assays

Four proteins identified by the iTRAQ analysis were measured by enzyme-linked immunoabsorbent assay (ELISA) from non-pooled samples to validate the iTRAQ findings.

These proteins were alpha-2-macroglobulin (A2M), retinol binding protein 4 (RBP4), serum amyloid A1 (SAA1) and apolipoprotein A1 (ApoA1). They were selected for their biological relevance and differential abundance between AIS C improvers and non-improvers, implying potential as biomarkers of neurological outcome prediction. A2M, RBP4 and SAA1 were assessed using a human DuoSet® ELISAs (R&D Systems, Abingdon, UK). ApoA1 was assessed using a human Quantikine® ELISA (R&D Systems, Abingdon, UK). Samples were diluted 1:600,000 for A2M and RBP4, 1:100 for SAA1 and 1:20,000 for ApoA1 in the respective assay kit diluent. Samples that were above the assay detection limit were rerun at 1:300 and 1:40,000 for SAA1 and ApoA1 respectively. All ELISAs were carried out according to the manufacturer’s protocol. Protein concentrations were normalised to the sample dilution factor. Statistical analysis was performed using the statistical programming language R version 4.2.1 (2022-06-23). Pairwise t tests with bonferroni adjusted P-values with the R rstatix package were used to assess differential abundance.

## 4. Results

Plasma from American Spinal Injury Association (ASIA) grade C SCI patients (total n=17) contrasting those who experienced an AISA grade conversion (n=10), and those who did not (n=7) collected within 2 weeks, and at approximately 3 months post-injury (Improvers n=9 vs Non-improvers n=6). Relative protein abundance in AIS grade A (n=10) and grade D (n=11) patients was also examined.

In the interest of brevity, only the plots of acute and subactue AIS C improvers VS non-improvers are included here, please see the supplemental data for the other comparisons (section Section 6).

### 4.1. iTRAQ analyses

### 4.2. Differential protein abundances

AIS C improvers had 18 more abundant proteins and 49 less abundant proteins at the acute phase relative to non-improvers. Similarly, at the subacute phase, AIS C improvers had 34 more abundant proteins and 34 less abundant proteins relative to non-improvers. The AIS A group had 56 more abundant and 9 less abundant proteins respectively relative to non-improvers. Acutely, AIS C improvers relative to AIS A and D had 21 and 53 more abundant and 46 and 12 less abundant proteins.

Please see the Tables S1, S2 & S3 for a full list of protein fold change changes.

### 4.3. Heatmaps

The majority of the pathways associated with the proteins identified by these iTRAQ experiments are related to the complement cascade and platelet activity (Figures 1, 2). There are also several pathways implicated in metabolic processes, particularly with apolipoproteins and retinoids.

**Figure 1:**
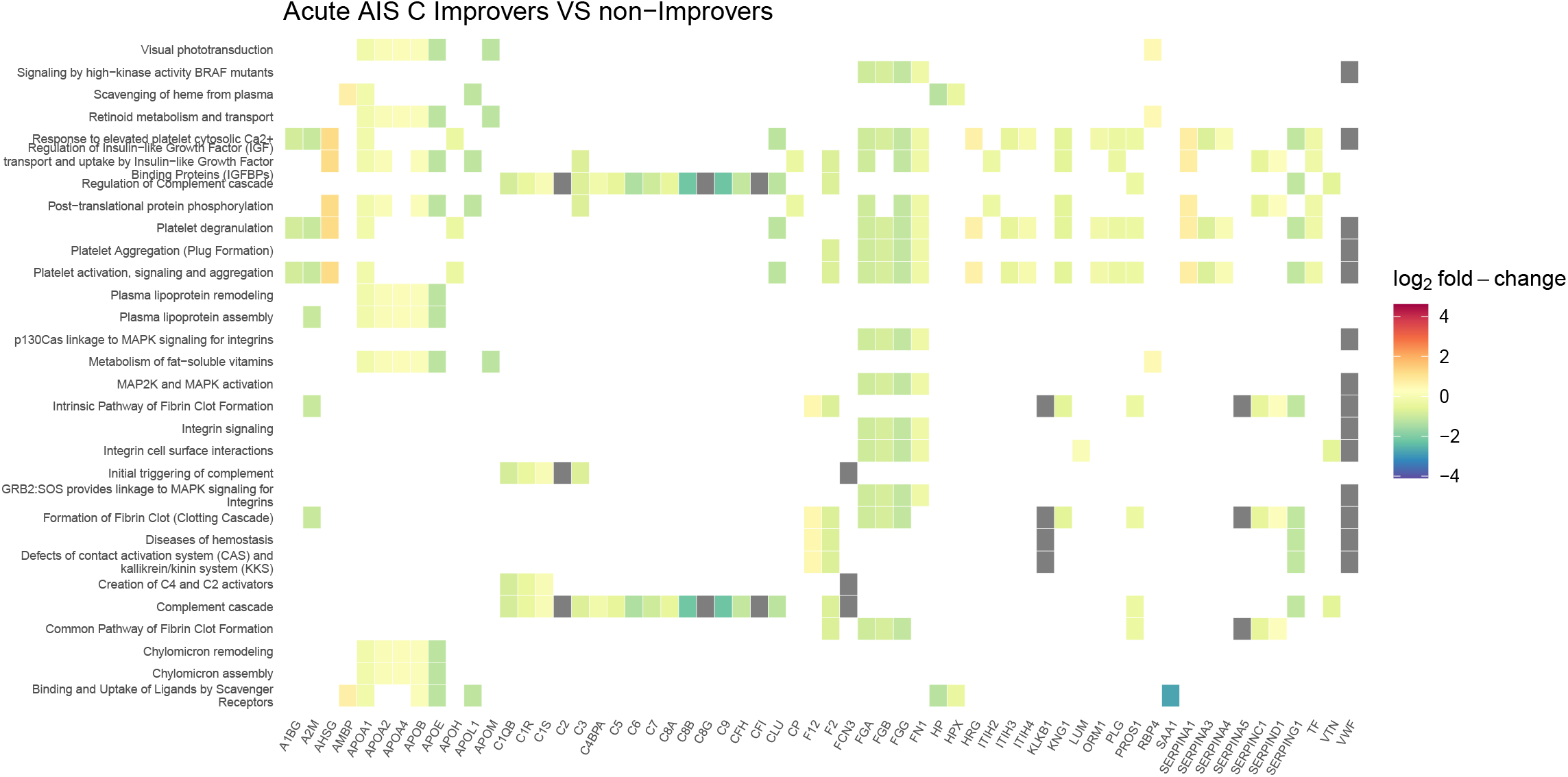
Heatmap denoting the log_2_ fold change of proteins in plasma collected 2-weeks post-injury, and the biological pathways these proteins are associated with on Reactome. This compares AIS C SCI patients who experienced an AIS grade improvement and those who did not.

**Figure 2:**
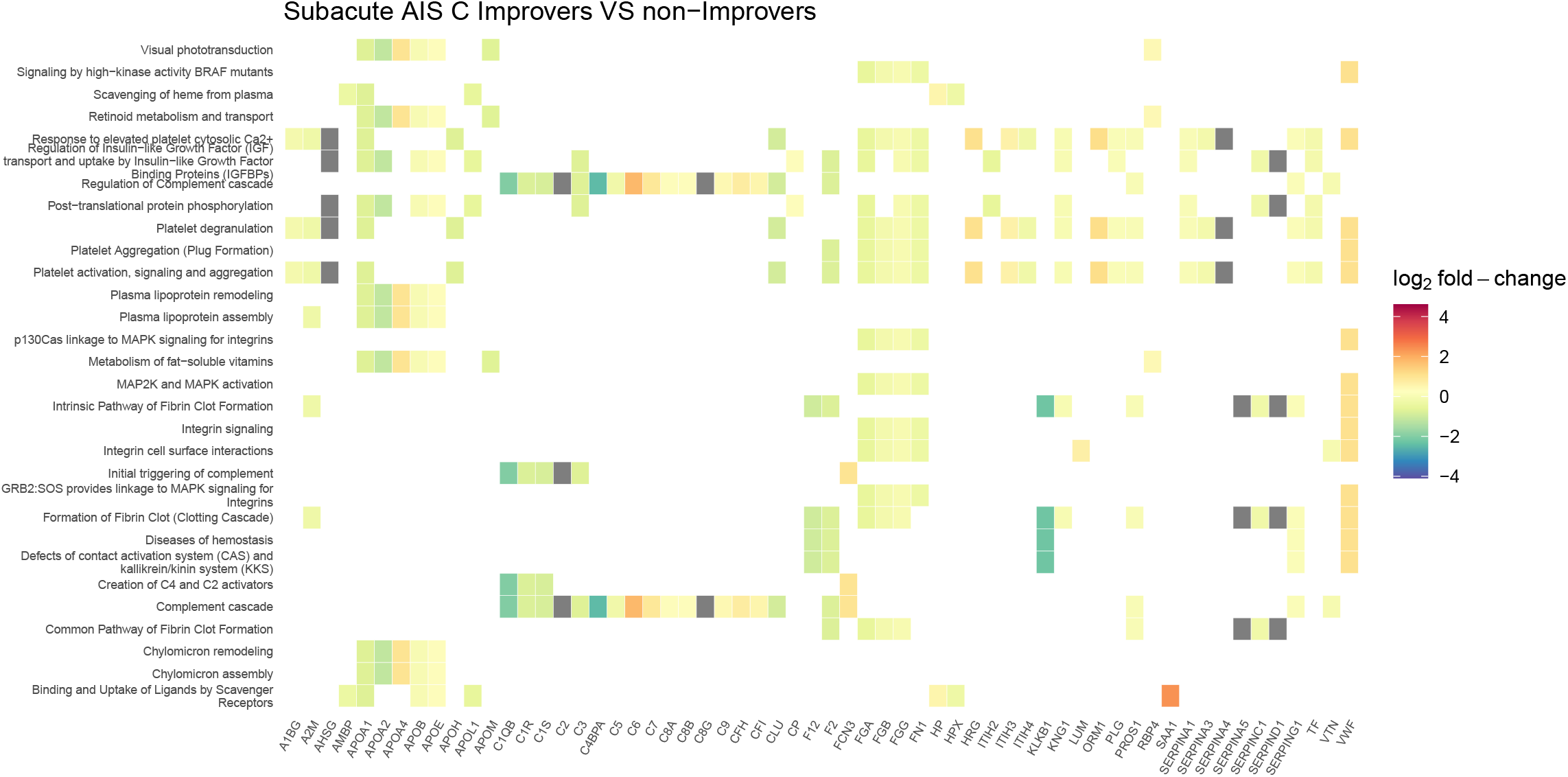
Heatmap denoting the log_2_ fold change of proteins in plasma collected 3-months post-injury, and the biological pathways these proteins are associated with on Reactome. This compares AIS C SCI patients who experienced an AIS grade improvement and those who did not.

Similarly to the iTRAQ data, many of the Reactome pathways are associated with the complement cascade and platelets activation (Figures 3, 4).

**Figure 3:**
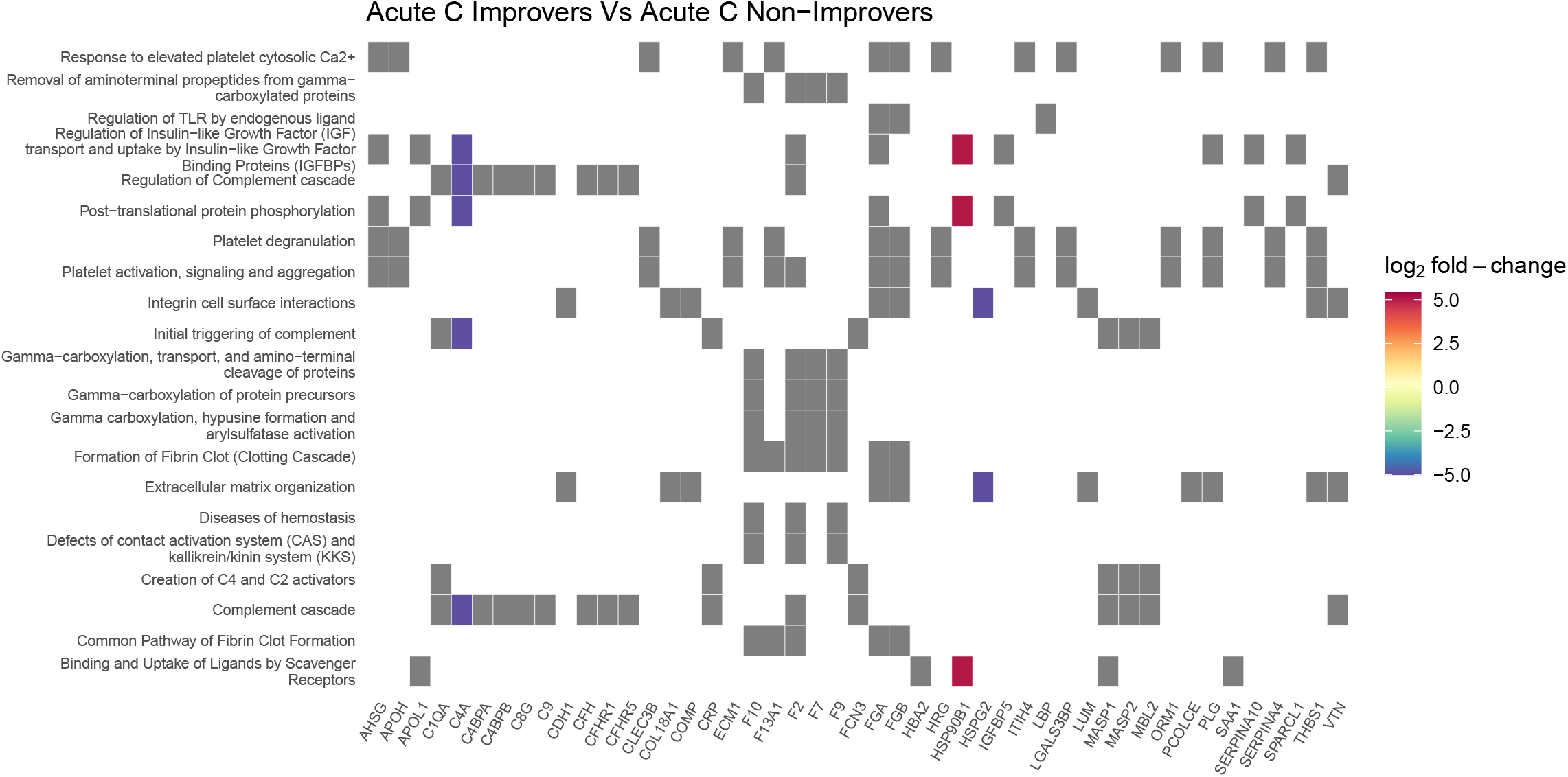
Heatmap denoting the log_2_ fold change of proteins in plasma collected 2-weeks post-injury, and the biological pathways these proteins are associated with on Reactome. This compares AIS C SCI patients who experienced an AIS grade improvement and those who did not. Grey blocks denote proteins not present in the comparison.

**Figure 4:**
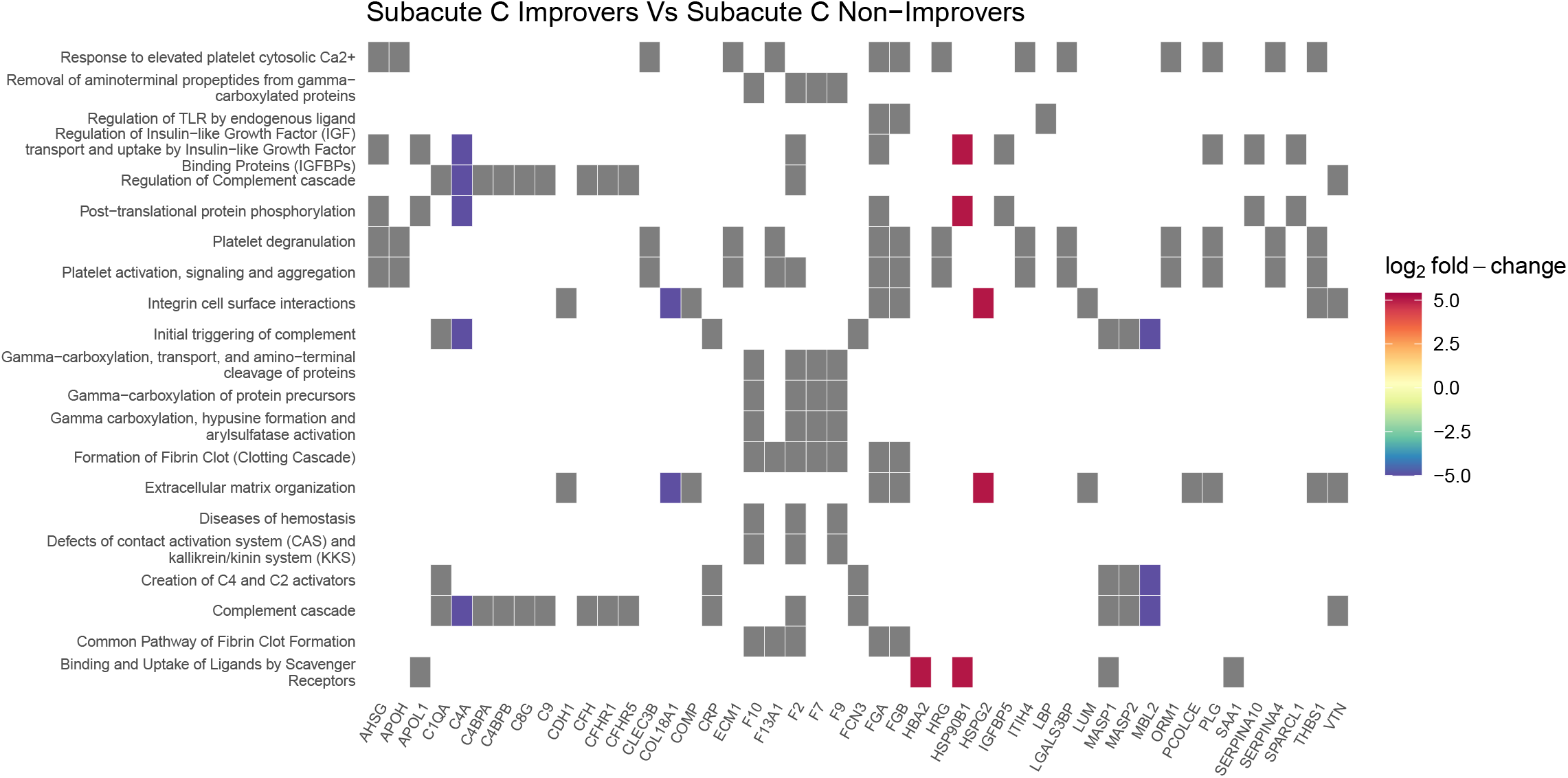
Heatmap denoting the log_2_ fold change of proteins in plasma collected 3-months post-injury, and the biological pathways these proteins are associated with on Reactome. This compares AIS C SCI patients who experienced an AIS grade improvement and those who did not. Grey blocks denote proteins not present in the comparison.

### 4.4. Network analysis of Differentially Abundant Proteins between AIS C improvers and non-improvers

Similar to the heatmaps, network plots highlighted that the majority of proteins changes were associated with the complement cascade and pathways linked to platelet activity (Figures 5, 6). Several proteins were also associated with the regulation of insulin-like growth factor.

**Figure 5:**
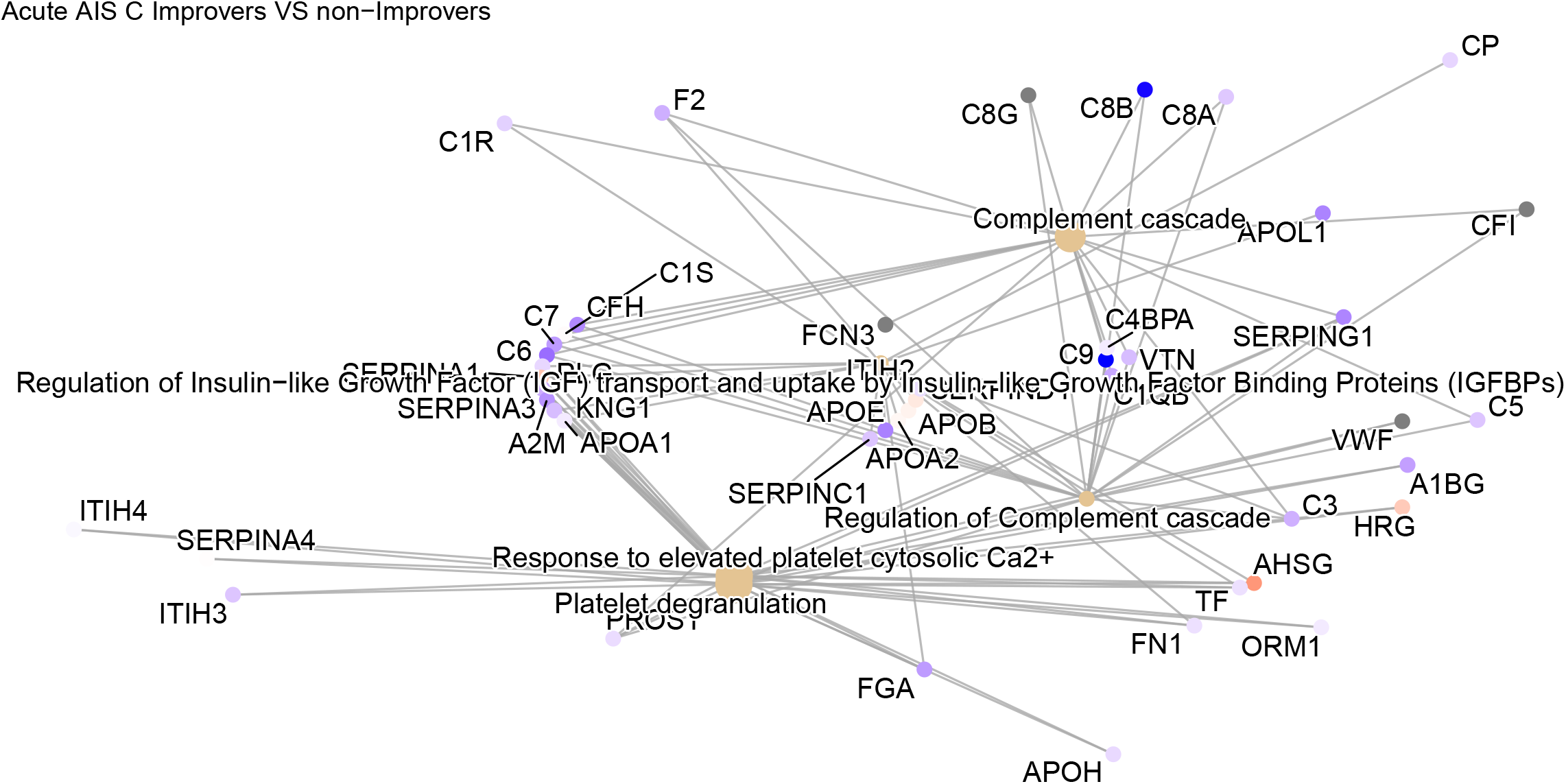
Network plot denoting the log_2_ fold change of proteins in plasma collected 2-weeks post-injury, and the biological pathways these proteins are associated with on Reactome. This compares AIS C SCI patients who experienced an AIS grade improvement and those who did not.

**Figure 6:**
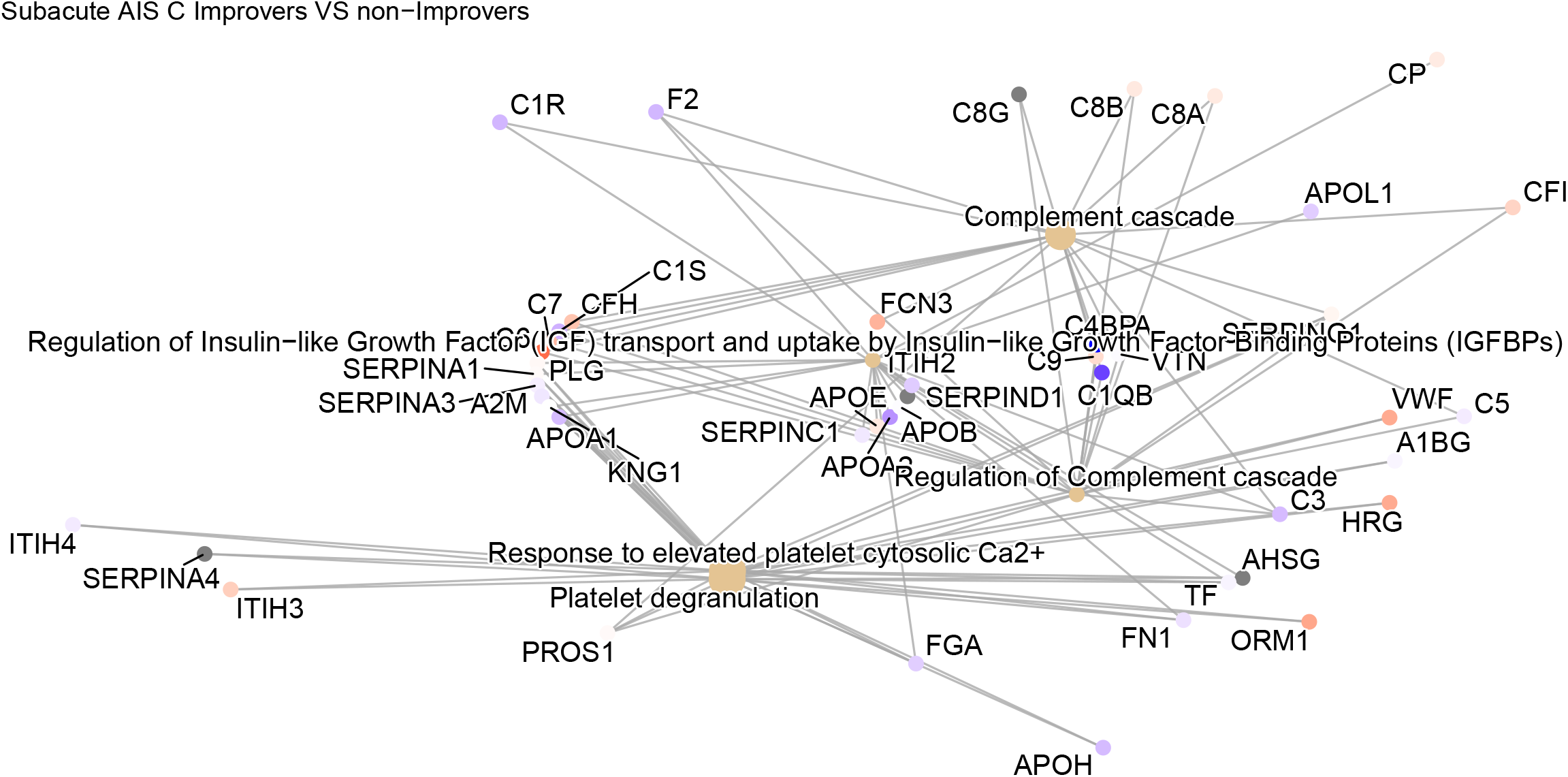
Network plot denoting the log_2_ fold change of proteins in plasma collected 3-months post-injury, and the biological pathways these proteins are associated with on Reactome. This compares AIS C SCI patients who experienced an AIS grade improvement and those who did not.

Similarly to the heatmaps and the iTRAQ data, network plots derived using the label-free data highlight the majority of differential proteins are associated with the complement cascade and pathways linked to platelets (Figures 7, 8).

**Figure 7:**
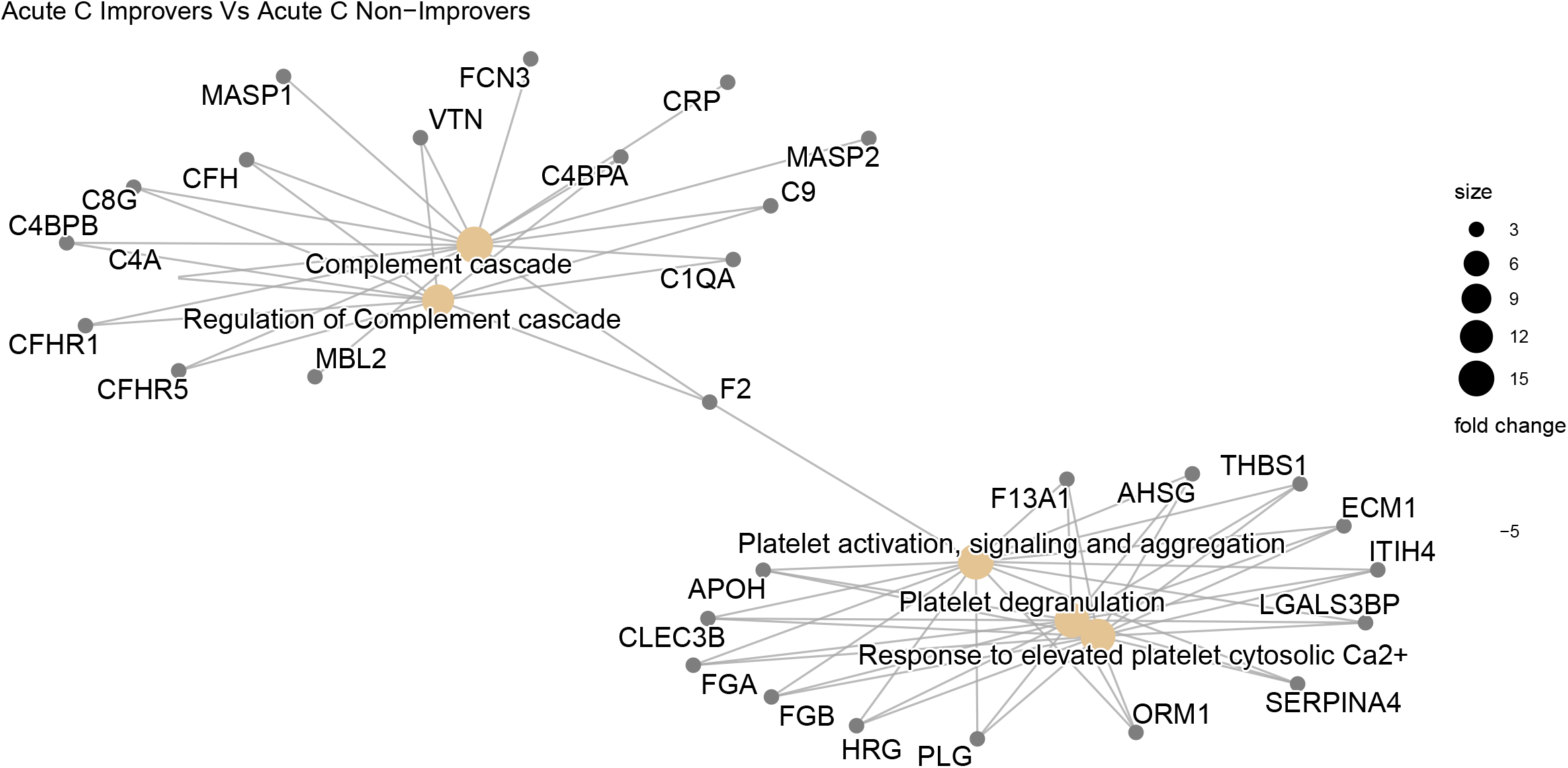
Network plot denoting the log_2_ fold change of proteins in plasma collected 2-weeks post-injury, and the biological pathways these proteins are associated with on Reactome. This compares AIS C SCI patients who experienced an AIS grade improvement and those who did not.

**Figure 8:**
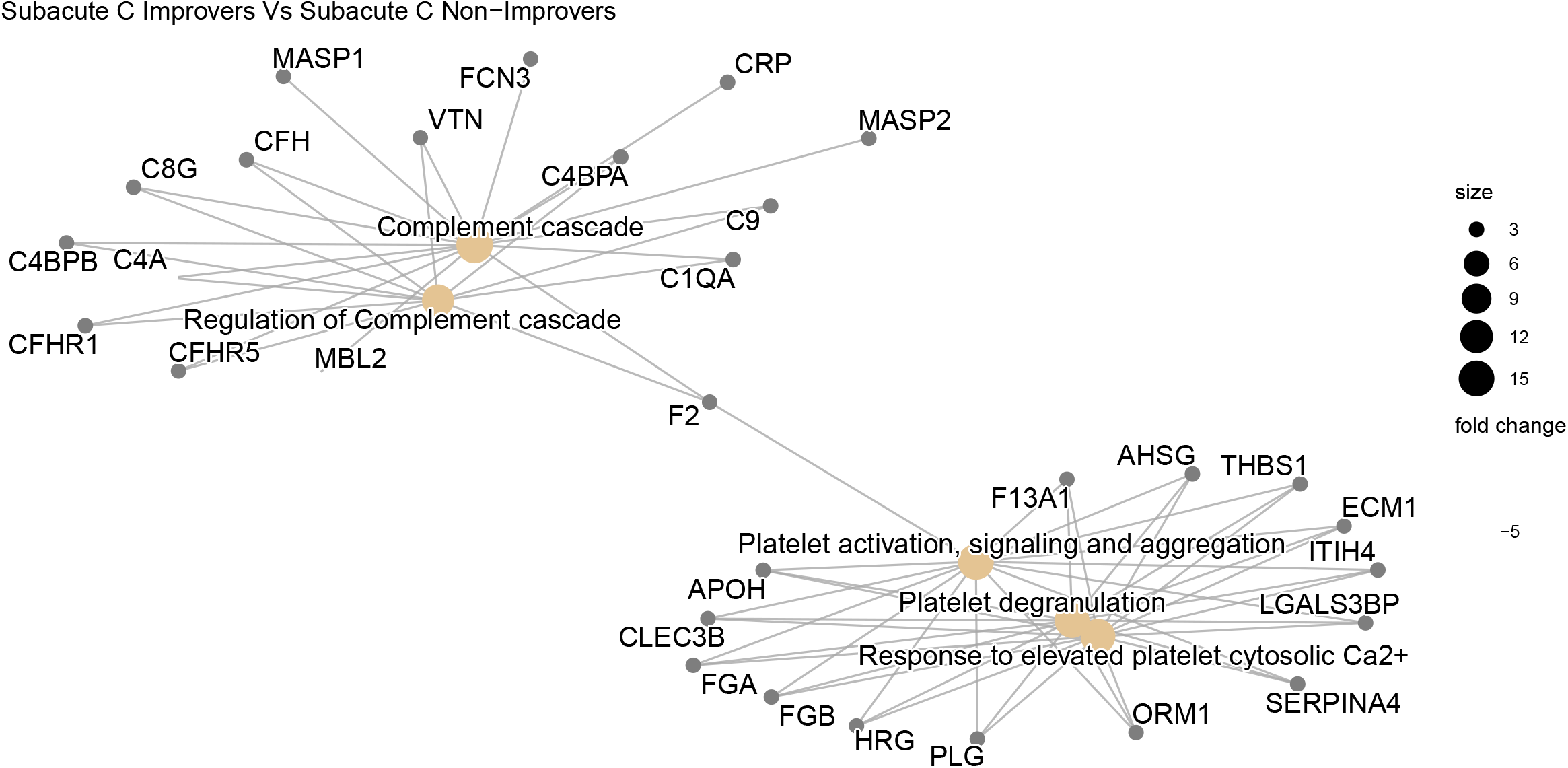
Network plot denoting the log_2_ fold change of proteins in plasma collected 3-months post-injury, and the biological pathways these proteins are associated with on Reactome. This compares AIS C SCI patients who experienced an AIS grade improvement and those who did not.

### 4.5. Pathway analysis of Differentially Abundant Proteins between AIS C improvers and non-improvers

Pathway analysis via the `pathview` R package returned the complement and coagulation cascade to be on the sole significant KEGG pathway to derive from the OpenMS analysed data. The majority of the proteins present in this pathway were less abundant in the 2-week post-injury plasma of AIS C patients who experienced an AIS grade conversion and those who did not (Figure 9).

**Figure 9:**
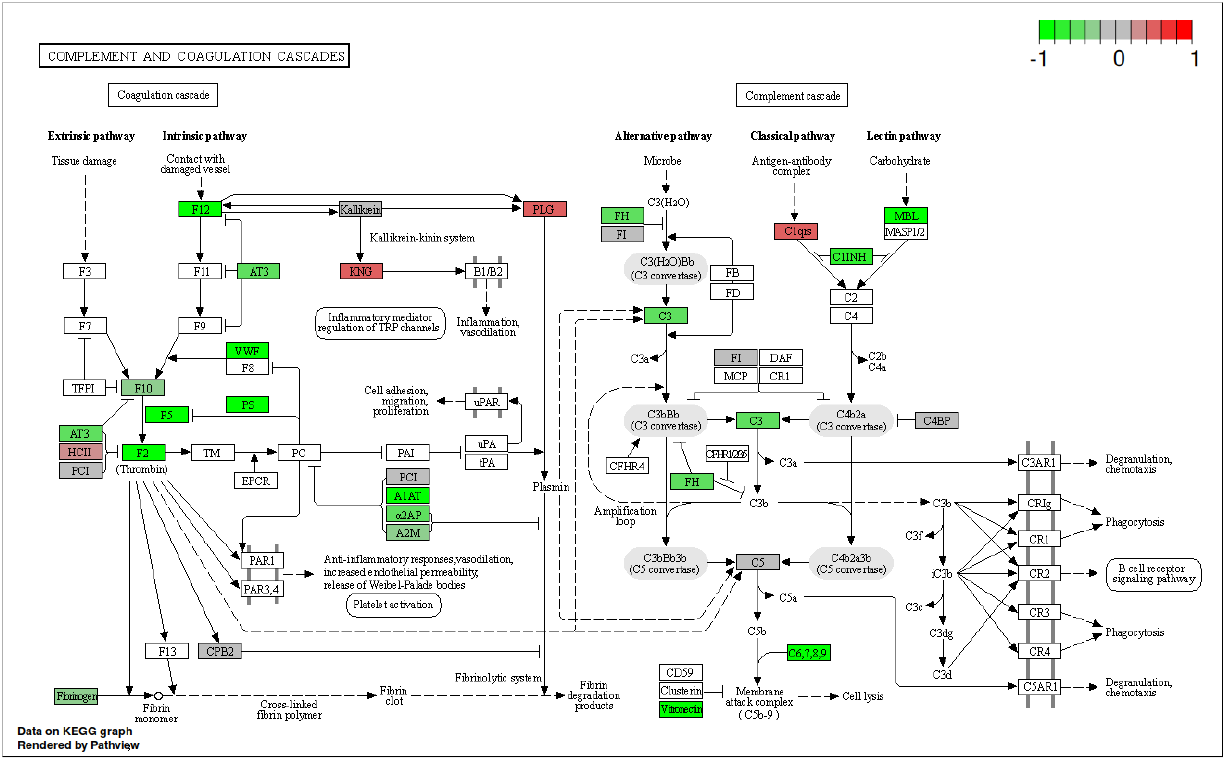
KEGG complement cascade pathway annotated with log_2_ fold change of proteins in plasma collected 2-weeks post-injury. This compares AIS C SCI patients who experienced an AIS grade improvement and those who did not.

Similarly to the iTRAQ pathway analysis, the label free data analysed via the pathview R package returned the complement and coagulation cascade to be the sole significant KEGG pathway derived from the OpenMS analysed data. The majority of the proteins present in this pathway were less abundant 2-weeks post-injury in the plasma of AIS C patients who experienced an AIS grade conversion than those who did not (Figure 10).

**Figure 10:**
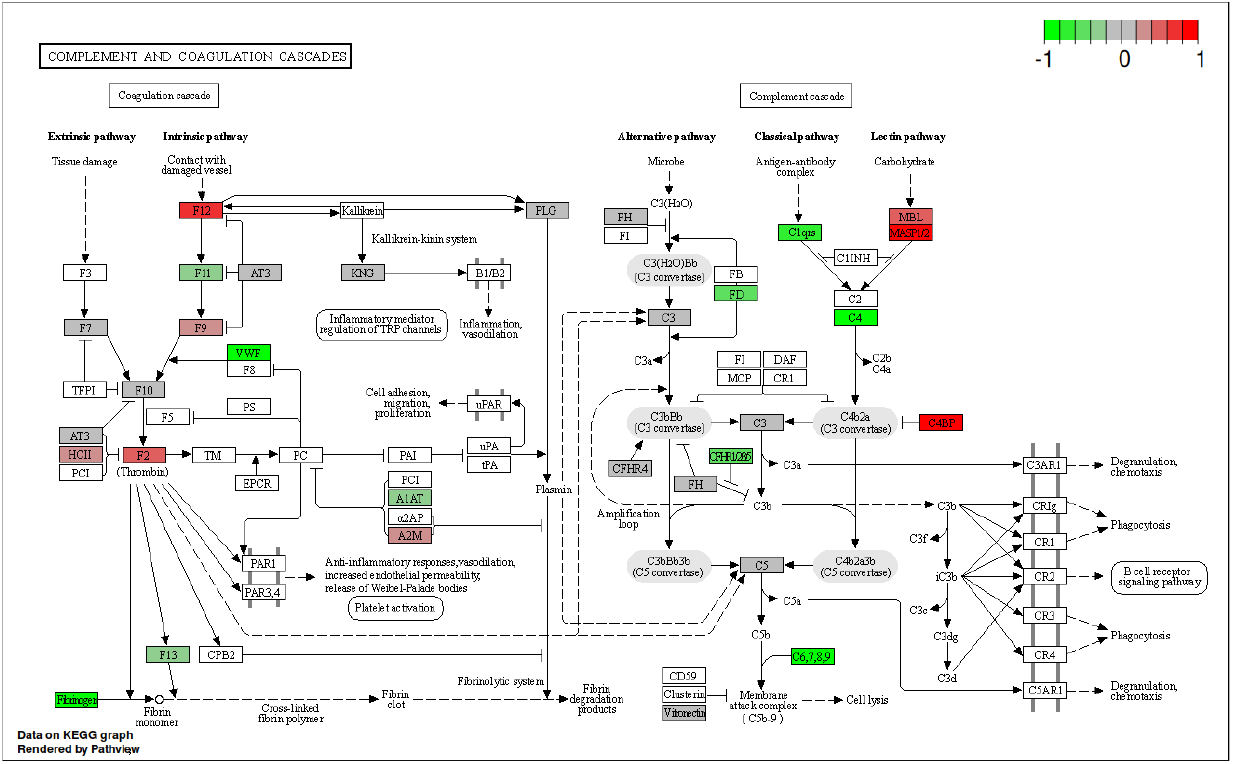
KEGG complement cascade pathway annotated with log_2_ fold change of proteins in plasma collected 2-weeks post-injury. This compares AIS C SCI patients who experienced an AIS grade improvement and those who did not.

### 4.6. Validation of Proteomic Data using ELISA

No statistically significant difference between groups for A2M abundance in plasma via DuoSet® ELISAs, though there were outliers in the AIS A and D groups, and particularly in the AIS C patients at 3-months who did not experience an AIS grade conversion (Figure 11).

**Figure 11:**
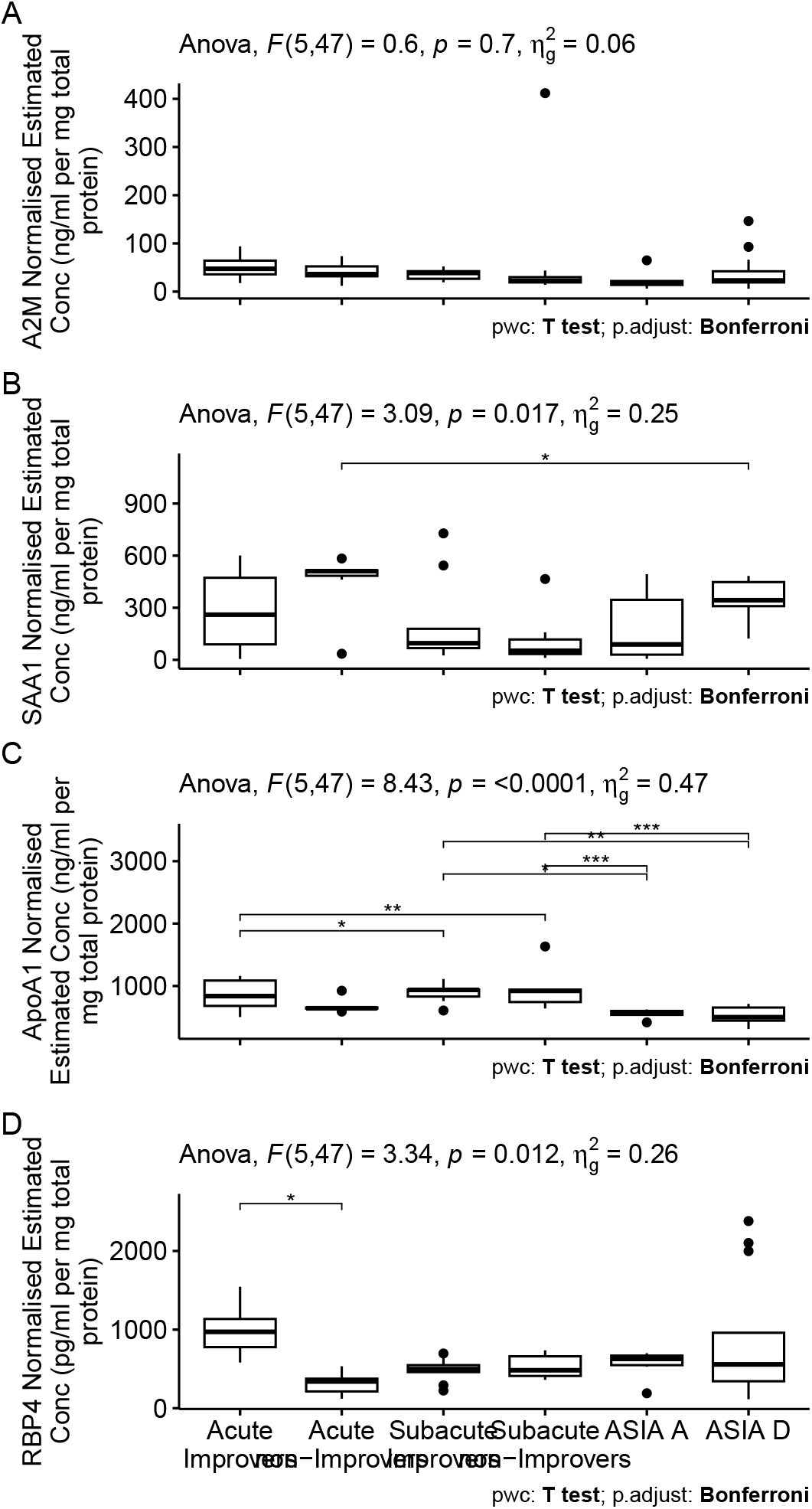
Normalised estimated concentration of *α*-2-macroglobulin (A), serum amyloid A1 (B), apolipoprotein A1 (C) and retinol binding protein 4 (D). Estimates were calculated from the optical density of a standard curve produced via a DuoSet® ELISA. Plasma from each patient that made up the pooled iTRAQ samples was assayed and pairwise t-tests with bonferroni adjusted P-values were performed to assess differential abundance.

A significant difference was found between AIS C non-improvers at 2-weeks and AIS D for SAA1, with outliers in AIS C non-improvers at 2-weeks, and both AIS C improvers and non-improvers at 3-months post-injury (Figure 11). For ApoA1 plasma abundance estimated via Quantikine® ELISAs, statistically significant differences were found between AIS C improvers at 2-weeks and both AIS C improvers and non-improvers at 3-months, AIS C 3-month improvers and AIS A and D, and AIS C 3-month non-improvers and AIS A and D (Figure 11). A statistically significant difference was also found between AIS C improvers and non-improvers at 2-weeks post-injury for RBP4 (Figure 11).

### 4.7. Comparing iTRAQ and label-free proteins

A total of 87 and 79 unique proteins were identified across the label-free and iTRAQ experiments respectively, with a modest overlap of 26 proteins found using both techniques.

## 5. Discussion

This is the first study, to our knowledge, to investigate the plasma proteome in SCI patients whose AIS scores either improved or did not improve post injury and also to compare these to AIS grade A and D patients. We have used two proteomic techniques allowing us to profile both high and low abundance proteins, in order to identify protein candidate biomarkers which may have potential to predict neurological improvement within the acute setting. Moreover, this data can better inform us of the biology underlying neurological improvement or stability in a cohort of patients being conservatively managed post SCI.

Briefly, for processing of proteomic data, we compared the performance of the mass spectrometry vendor (ABSciex) provided ProteinPilot (version 4.5) and OpenMS (version 2.6.0). As there were only modest difference in both the proteins identified and the respective fold changes (data not shown), we opted to use OpenMS for the greater transparency and reproducibility it offers as open-source software.

This study has highlighted a number of proteins that may be able to discriminate in, the acute phase following injury, between AIS grade C patients who either improve or do not improve by an AIS grade following SCI. The most promising of these is Retinol Binding Protein 4 (RBP4) which was demonstrated to be increased in improvers compared to non-improvers in the acute phase. Further this change could be confirmed using ELISA, which may provide a more clinically useful means of assessment on a wide scale.

RBP4 is synthesised in the liver and binds retinol that is released following vitamin A deficiency. ^31^ Once delivered to target cells, retinol can either be converted to retinaldehyde, which is required for functional vision, or oxidised to retinoic acid, which is a ligand for nuclear receptors, thus regulating gene expression. ^32,33^ The role of retinoid signalling in spinal cord and motor neuron differentiation, including development of regions of the spinal cord has been outlined, and implies a possible involvement in maintaining motor neuron integrity. ^34,35^ The mRNA of a rodent homologue of RBP was found to be up-regulated at 24 hours post-SCI and may promote cell proliferation and regeneration by increasing retinoid metabolism. ^36,37^

Another study of amyotrophic lateral sclerosis (ALS), a neurodegenerative disease, comparing gene expression between post-mortem spinal cord samples of ALS and controls similarly observed up-regulation of RBP1 in ALS spinal cord. ^38^ Furthermore, a transgenic mouse study reported retinoid signalling may contribute to the retained plasticity and regenerative potential of the mature spinal cord. ^39^ Collectively, these results might support a hypothesis that AIS C improvers have increased levels of RBP4 and this relates to improved capacity for neuronal regeneration/plasticity. Whether this is due to increased expression or due to higher vitamin A dietary intake is unclear from this data.

Alongside RBP4, a number of other protein abundance differences across the different biological comparisons were identified in proteins associated with liver function. Our previous work investigating the potential of routinely measured haematological analytes for predicting neurological outcome in SCI patients also highlighted several proteins that were linked with liver function; thus providing further support to the theory that liver status is relevant to differential functional recover. ^17,16^ The pathway analysis specifically indicated that the acute phase response (APR) is implicated.

The APR is the bodies first response to infection or injury, including SCI. This systemic response is largely coordinated by factors released from the liver, but the APRs effects extend to multiple peripheral organs including the kidneys, lungs and spleen. ^40,41,42,4^ This hepatic response is typically transient and quickly fades, but prolonged liver inflammation and pathology has been observed in rodent SCI models. ^43,44^ Basic liver functions are chronically impaired by SCI, including metabolising carbohydrates, fats and proteins, storage of minerals vitamins and glycogen and filtering blood from the digestive tract. ^45,46,47,48,44^

The acute (1-7 days) liver response to SCI is well documented; the inflammatory cytokines including TNF*α*, IL-1*α*, IL-1*β* and IL-6, released at the injury site, reach the liver through the bloodstream. ^42,49^ This provokes the liver to enter the APR and produce acute phase proteins thus stimulating a greater immune response. ^50,42^ The hepatocytes that make up the majority of the liver biomass, express receptors that bind the aforementioned inflammatory cytokines; similarly the hepatic macrophage Kupffer cells also bind these cytokines, complement proteins and lipopolysaccharide (LPS) and swiftly remove microorganisms, endotoxins and other debris from the blood. ^51,52,53,54^ Furthermore, it has been suggested that liver inflammation and Kupffer cells activity promote recruitment of leukocytes to the injury site in brain or spinal trauma, potentially enhancing CNS injury.^50,54^ For example, a rodent study demonstrated depletion of Kupffer cells prior to injury resulted in few neutrophils infiltrating the injury site. ^41,55^

Another protein that out label-free proteomic data highlights is Peroxiredoxin 2 (PRX-2), which was detected acutely in AIS C improvers and AIS D patients, and subacutely in AIS A and AIS D. Peroxiredoxins are a large and highly conserved family of enzymes that reduce peroxides. PRX-2 is highly abundant in RBCs and intracellularly serves as an important anti-oxidant role in various cells types, including neurons. ^56^ By contrast, extracellular PRX-2 has been suggested to act as an inflammatory DAMP, leading microglia and macrophages to release a plethora of pro-inflammatory factors. ^57,58,59^

An *in vitro* primary neurons and microglia co-culture study reported PRX-2 activating microglia via TLR-4, potentially leading to neuronal apoptosis. ^60^ A murine study found over-expression of PRX-2 attenuated oxidative stress and neuronal apoptosis following subarachnoid haemorrhage. ^61^ Over-expression of PRX-2 is speculated to protect again ischaemic neuronal injury by modulating the redox-sensitive thioredoxin-apoptosis signal-regulating kinase (ASK) 1 signalling complex.^62^ Several molecular chaperones can interact with ASK1, including thioredoxin and TNF receptor-associated factor 6. ^63^ The dissociation of the thioredoxin-ASK1 complex activates ASK1. PRX-2 is oxidised after scavenging free radicals, whereupon its antioxidantive activity is reduced. This inactivation can be reversed by the thioredoxin-thioredoxin reductase system, whereby oxidised PRX-2 can regain its activity by reducing thioredoxin, leading to the dissociation of the thioredoxin-ASK1 complex. ^64^ Additionally, oxidised PRX-1 can inhibit ASK1-induced apoptosis via the thioredoxin-binding domain on ASK1. ^65^

The presence of PRX-2 in acute AIS C improvers and absence in acute C non-improvers suggests the protein could indicate a more protective action against oxidative stress, and implies the protein has potential value as a biomarker of functional outcomes. Similarly, PRX-2 may be acting as a healthy response to trauma-induced oxidative stress in both acute AIS D, although the persistence to the subacute time-point is less clear. Likewise, the presence of PRX-2 in AIS A subacutely, but not acutely is more perplexing. It should be noted that as plasma was used and cells were lysed, there is no distinction between intracellular and extracellular PRX-2 in this data. Perhaps in the more severe AIS A injury, secondary injuries, including oxidative stress, are greater and so persist to the subacute time-point. The acute absence may be a result of an overwhelmed physiology unable to respond or prioritise managing oxidative stress.

Pathway analysis from both the iTRAQ and label-free experiments identified the complement and coagulation cascades as a significant pathway of interest. More broadly, the trend in this data is for proteins in the complement pathway is lower abundance, or inhibitory proteins such as C4BP to be more abundant, in the acute improvers. C3 for instance, cleavage of which is vital for complement activation, was less abundant in acute AIS C improvers relative to non-improvers. This finding is in line with a genetic C3 knockout study in mice which reported better neurological scores 2 days post-injury, reduced residual consolidated neurological deficit at 21 days and display minor change in reduced gliosis (20% decrease at 1h timepoint) but a three-to-fourfold decrease in neutrophil infiltration, resulting in enhanced regeneration of axons. ^66^ Another study using a similar C3 knockout model reported improved neurological scores at acute and long-term time points. ^67^ These results imply that the complement cascade is a particularly important component of a differential response to neurological injury which ultimately leads to greater functional recovery. Given the complexity of the complement cascade and the limited time points in this study, further work is needed to elucidate which facets of the cascade are outcome modifying, and at which stages post-injury.

AIS A and D samples were included largely to compare to the AIS C improvers. If the improvers were all just a less severe AIS C, we might expect them to be more similar to the AIS D samples, with the non-improvers being closer to the As. As this is not what we observed, we can conclude there is more to the differential functional recovery than initial injury severity.

The small number of statistically significant proteins speaks to the variability of human plasma samples, and is likely exacerbated by the inconstant timing of sample collection relative to injury. Thus, a repeat of this experiment with a larger sample size will likely reveal many more proteins of potential interest. Regardless, this study has highlighted RPB4 and PRX-2 as potential biomarkers of functional recover. We have also highlighted the complement cascade as being a particularly important pathway in differential recovery. Additional investigation of these proteins, but also the complement cascade more broadly, particularly at more acute time points, would also be valuable. Furthermore, a metabolomic analysis with a similar samples would greatly compliment this work, particularly with regards to investigating further links to the livers role in neurological recovery. Similarly, this additional work could complement the AIS A and D sample data, which may reveal further insights from this data.

## 6. Supplementary material

### 6.1. Session Information

**R version 4.2.1 (2022-06-23)**

**Platform:** aarch64-apple-darwin20 (64-bit)

8

**locale:** en_US.UTF-8||en_US.UTF-8||en_US.UTF-8||C||en_US.UTF-8||en_US.UTF-

**attached base packages:**

- stats
- graphics
- grDevices
- datasets
- utils
- methods
- base

**other attached packages:**

- MSstats(v.4.6.2)
- STRINGdb(v.2.10.1)
- ReactomePA(v.1.42.0)
- huxtable(v.5.5.2)
- pander(v.0.6.5)
- labelled(v.2.10.0)
- gtsummary(v.1.7.0)
- gt(v.0.8.0)
- rlang(v.1.0.6)
- bookdown(v.0.27)
- lime(v.0.5.2)
- RColorBrewer(v.1.1-3)
- ggVennDiagram(v.1.2.0)
- DiagrammeR(v.1.0.9)
- lubridate(v.1.8.0)
- patchwork(v.1.1.1)
- cowplot(v.1.1.1)
- readxl(v.1.4.0)
- BiocManager(v.1.30.18)
- knitr(v.1.39)
- rmarkdown(v.2.14)
- data.table(v.1.14.2)
- naniar(v.0.6.1)
- psych(v.2.2.5)
- Hmisc(v.4.7-0)
- Formula(v.1.2-4)
- survival(v.3.5-3)
- lattice(v.0.20-45)
- bibtex(v.0.4.2.3)
- captioner(v.2.2.3)
- forcats(v.0.5.2)
- stringr(v.1.4.0)
- dplyr(v.1.0.10)
- purrr(v.0.3.4)
- readr(v.2.1.3)
- tidyr(v.1.2.1)
- tibble(v.3.1.8)
- ggplot2(v.3.4.0)
- tidyverse(v.1.3.1)
- kableExtra(v.1.3.4)

**loaded via a namespace (and not attached):**

- utf8(v.1.2.2)
- proto(v.1.0.0)
- lme4(v.1.1-31)
- tidyselect(v.1.2.0)
- RSQLite(v.2.2.20)
- AnnotationDbi(v.1.60.0)
- htmlwidgets(v.1.6.1)
- grid(v.4.2.1)
- BiocParallel(v.1.32.5)
- scatterpie(v.0.1.8)
- munsell(v.0.5.0)
- preprocessCore(v.1.60.1)
- codetools(v.0.2-19)
- chron(v.2.3-58)
- interp(v.1.1-3)
- withr(v.2.5.0)
- colorspace(v.2.0-3)
- GOSemSim(v.2.24.0)
- Biobase(v.2.58.0)
- rstudioapi(v.0.14)
- stats4(v.4.2.1)
- DOSE(v.3.24.2)
- labeling(v.0.4.2)
- GenomeInfoDbData(v.1.2.9)
- mnormt(v.2.1.1)
- polyclip(v.1.10-4)
- bit64(v.4.0.5)
- farver(v.2.1.1)
- treeio(v.1.23.0)
- vctrs(v.0.5.1)
- generics(v.0.1.3)
- gson(v.0.0.9)
- xfun(v.0.36)
- R6(v.2.5.1)
- GenomeInfoDb(v.1.34.6)
- graphlayouts(v.0.8.4)
- RVenn(v.1.1.0)
- bitops(v.1.0-7)
- cachem(v.1.0.6)
- fgsea(v.1.24.0)
- gridGraphics(v.0.5-1)
- assertthat(v.0.2.1)
- scales(v.1.2.1)
- ggraph(v.2.1.0)
- nnet(v.7.3-18)
- enrichplot(v.1.18.3)
- gtable(v.0.3.1)
- log4r(v.0.4.3)
- tidygraph(v.1.2.2)
- systemfonts(v.1.0.4)
- splines(v.4.2.1)
- lazyeval(v.0.2.2)
- broom(v.1.0.2)
- checkmate(v.2.1.0)
- yaml(v.2.3.6)
- reshape2(v.1.4.4)
- modelr(v.0.1.10)
- backports(v.1.4.1)
- qvalue(v.2.30.0)
- tools(v.4.2.1)
- ggplotify(v.0.1.0)
- gplots(v.3.1.3)
- ellipsis(v.0.3.2)
- BiocGenerics(v.0.44.0)
- gsubfn(v.0.7)
- hash(v.2.2.6.2)
- Rcpp(v.1.0.9)
- plyr(v.1.8.8)
- base64enc(v.0.1-3)
- visNetwork(v.2.1.2)
- zlibbioc(v.1.44.0)
- RCurl(v.1.98-1.9)
- sqldf(v.0.4-11)
- rpart(v.4.1.19)
- deldir(v.1.0-6)
- viridis(v.0.6.2)
- S4Vectors(v.0.36.1)
- haven(v.2.5.1)
- ggrepel(v.0.9.2)
- cluster(v.2.1.4)
- fs(v.1.5.2)
- magrittr(v.2.0.3)
- reactome.db(v.1.82.0)
- reprex(v.2.0.2)
- ggnewscale(v.0.4.8)
- hms(v.1.1.2)
- evaluate(v.0.20)
- HDO.db(v.0.99.1)
- jpeg(v.0.1-10)
- IRanges(v.2.32.0)
- gridExtra(v.2.3)
- shape(v.1.4.6)
- compiler(v.4.2.1)
- KernSmooth(v.2.23-20)
- shadowtext(v.0.1.2)
- crayon(v.1.5.2)
- minqa(v.1.2.5)
- htmltools(v.0.5.4)
- ggfun(v.0.0.9)
- tzdb(v.0.3.0)
- aplot(v.0.1.9)
- visdat(v.0.5.3)
- DBI(v.1.1.3)
- tweenr(v.2.0.2)
- dbplyr(v.2.3.0)
- rappdirs(v.0.3.3)
- MASS(v.7.3-58.2)
- broom.helpers(v.1.11.0)
- boot(v.1.3-28.1)
- Matrix(v.1.5-3)
- cli(v.3.6.0)
- MSstatsConvert(v.1.8.2)
- marray(v.1.76.0)
- parallel(v.4.2.1)
- gower(v.1.0.1)
- igraph(v.1.3.5)
- pkgconfig(v.2.0.3)
- foreign(v.0.8-84)
- xml2(v.1.3.3)
- foreach(v.1.5.2)
- ggtree(v.3.6.2)
- svglite(v.2.1.1)
- webshot(v.0.5.4)
- XVector(v.0.38.0)
- rvest(v.1.0.3)
- snakecase(v.0.11.0)
- yulab.utils(v.0.0.6)
- digest(v.0.6.31)
- janitor(v.2.1.0)
- graph(v.1.76.0)
- Biostrings(v.2.66.0)
- cellranger(v.1.1.0)
- fastmatch(v.1.1-3)
- tidytree(v.0.4.2)
- htmlTable(v.2.4.1)
- gtools(v.3.9.4)
- graphite(v.1.44.0)
- nloptr(v.2.0.3)
- lifecycle(v.1.0.3)
- nlme(v.3.1-162)
- jsonlite(v.1.8.4)
- limma(v.3.54.0)
- viridisLite(v.0.4.1)
- fansi(v.1.0.3)
- pillar(v.1.8.1)
- plotrix(v.3.8-2)
- KEGGREST(v.1.38.0)
- fastmap(v.1.1.0)
- httr(v.1.4.4)
- GO.db(v.3.16.0)
- glue(v.1.6.2)
- png(v.0.1-8)
- iterators(v.1.0.14)
- glmnet(v.4.1-6)
- bit(v.4.0.5)
- ggforce(v.0.4.1)
- stringi(v.1.7.12)
- blob(v.1.2.3)
- caTools(v.1.18.2)
- latticeExtra(v.0.6-30)
- memoise(v.2.0.1)
- renv(v.0.16.0)
- ape(v.5.6-2)

---

**Table S1:** OpenMS log_2_ fold changes in the plasma proteome of SCI patients from iTRAQ experiments. ‘Acute’ and ‘Subacute’ samples collected within 2 week and approximately 3-months post-injury respectively.

| Protein | Acute<br>AIS C<br>improvers<br>vs non-improvers | Subacute<br>AIS C<br>improvers<br>vs non-improvers | AIS C<br>improvers<br>acute vs<br>subacute | AIS C non-improvers<br>acute vs<br>subacute | AIS C<br>improvers<br>vs non-improvers | AIS A vs<br>D | AIS C im-<br>provers vs<br>A | AIS C im-<br>provers vs<br>D | AIS C non-improvers<br>vs A | AIS C non-improvers<br>vs D |
| --- | --- | --- | --- | --- | --- | --- | --- | --- | --- | --- |
| A1BG | -0.903182 | -0.101753 | -0.60878 | 0.192644 | 0.22527 | 0.79373 | -0.34976 | 0.44397 | -0.575028 | 0.218706 |
| A2M | -1.038579 | -0.246439 | -0.67606 | 0.116078 | -1.23010 | 1.42475 | -1.60298 | -0.17823 | -0.372883 | 1.051871 |
| AFM | -0.378848 | -1.224864 | 0.48152 | -0.364497 | 0.55179 | 1.19236 | -1.25661 | -0.06425 | -1.808399 | -0.616039 |
| AHSG | 1.179453 | — | -0.55453 | — | — | — | — | — | — | — |
| AMBP | 0.656200 | -0.343343 | 0.86066 | -0.138885 | -0.90233 | — | 1.20378 | — | 2.106113 | — |
| APCS | 0.149829 | 0.210894 | -0.01140 | 0.049664 | — | 0.35572 | — | — | -0.049457 | 0.306268 |
| APOA1 | -0.181674 | -0.692362 | -0.23376 | -0.744443 | -0.76773 | 0.69413 | -1.31728 | -0.62316 | -0.549553 | 0.144575 |
| APOA2 | 0.090014 | -1.146136 | -0.66676 | -1.902912 | — | — | — | — | — | — |
| APOA4 | 0.129596 | 0.963678 | -1.23128 | -0.397198 | -1.32541 | 0.78760 | -1.33467 | -0.54707 | -0.009263 | 0.778338 |
| APOB | 0.137923 | -0.016410 | -0.63328 | -0.787608 | -0.85704 | 0.52600 | -1.23459 | -0.70858 | -0.377547 | 0.148457 |
| APOE | -1.213375 | 0.293067 | -0.68845 | 0.817994 | -0.90783 | 0.77465 | -1.54775 | -0.77310 | -0.639919 | 0.134733 |
| APOH | -0.360239 | -0.702469 | -0.64449 | -0.986719 | -0.99966 | 2.81436 | -1.00918 | 1.80518 | -0.009516 | 2.804845 |
| APOL1 | -1.179076 | -0.519351 | -1.04403 | -0.384302 | -0.11528 | 0.56527 | 0.12993 | 0.69520 | 0.245210 | 0.810480 |
| APOM | -1.216797 | -0.681988 | 0.69348 | 1.228290 | — | 0.65618 | — | — | 0.666495 | 1.322676 |
| ATRN | — | — | -1.00630 | — | — | — | — | — | — | — |
| AZGP1 | 1.219168 | 1.025150 | 0.08114 | -0.112878 | -3.38895 | -3.64405 | 0.37029 | -3.27376 | 3.759240 | 0.115190 |
| C1QB | -0.841007 | -2.002039 | 0.70711 | -0.453921 | -1.97292 | 1.35633 | -2.00663 | -0.65030 | -0.033709 | 1.322622 |
| C1R | -0.433512 | -0.763216 | 0.03665 | -0.293055 | -0.14675 | 0.79761 | 0.35643 | 1.15404 | 0.503179 | 1.300786 |
| C1S | 0.029522 | -0.819374 | 0.16796 | -0.680940 | — | — | — | — | — | — |
| C2 | — | — | — | — | -2.55810 | 2.56410 | -2.59527 | -0.03117 | -0.037167 | 2.526930 |
| C3 | -0.744062 | -0.696858 | 0.06524 | 0.112441 | -1.07308 | 1.23884 | -2.16164 | -0.92280 | -1.088566 | 0.150276 |
| C4BPA | -0.181039 | -2.445498 | 1.66277 | -0.601693 | -1.23787 | 1.54897 | -1.84489 | -0.29592 | -0.607021 | 0.941952 |
| C5 | -0.544784 | -0.203123 | 0.92300 | 1.264662 | -0.72000 | 1.27105 | -1.67688 | -0.40583 | -0.956878 | 0.314172 |
| C6 | -1.393621 | 1.781702 | -1.30971 | 1.865613 | -3.04519 | 1.76424 | -3.25500 | -1.49076 | -0.209810 | 1.554427 |
| C7 | -0.964212 | 0.884808 | -0.78272 | 1.066304 | 0.99702 | 0.07087 | -1.11363 | -1.04277 | -2.110650 | -2.039785 |
| C8A | -0.511789 | 0.273656 | -0.76301 | 0.022431 | -2.81083 | 0.17312 | -2.12854 | -1.95541 | 0.682296 | 0.855420 |
| C8B | -2.195043 | 0.278904 | -1.59549 | 0.878459 | -1.89440 | -0.48026 | -0.95975 | -1.44001 | 0.934642 | 0.454381 |
| C8G | — | — | -1.63049 | — | — | — | — | — | — | — |
| C9 | -2.219906 | 0.453409 | -1.92498 | 0.748336 | -0.73459 | 0.64959 | -3.24243 | -2.59284 | -2.507839 | -1.858252 |
| CD5L | -0.929325 | -0.620473 | -0.71456 | -0.405706 | -2.46429 | 0.44825 | -2.32601 | -1.87776 | 0.138275 | 0.586529 |
| CFH | -1.123974 | 0.740695 | -1.64809 | 0.216580 | -1.03587 | 0.13801 | -1.32605 | -1.18804 | -0.290178 | -0.152168 |
| CFI | — | 0.535970 | — | 1.257811 | — | — | — | — | — | — |
| CLU | -1.195898 | -0.868185 | -0.17219 | 0.155521 | -1.36644 | 0.82520 | -2.19762 | -1.37242 | -0.831181 | -0.005985 |
| CP | -0.389206 | 0.256541 | -0.45373 | 0.192020 | -0.66575 | 0.42354 | -0.26958 | 0.15395 | 0.396174 | 0.819709 |
| F12 | 0.485201 | -0.939791 | 0.67029 | -0.754699 | -0.85343 | 0.55496 | -1.31458 | -0.75963 | -0.461154 | 0.093802 |
| F2 | -0.749308 | -0.756359 | 0.09829 | 0.091237 | -0.54088 | 1.16771 | -1.54762 | -0.37990 | -1.006738 | 0.160976 |
| FCN3 | — | 0.964478 | — | — | — | — | — | — | — | — |
| FGA | -0.959140 | -0.510905 | 0.48417 | 0.932405 | -1.01557 | 1.04867 | -1.47080 | -0.42212 | -0.455227 | 0.593445 |
| FGB | -0.833909 | -0.125377 | 0.06843 | 0.776960 | -0.83431 | 1.09511 | -1.46465 | -0.36955 | -0.630340 | 0.464768 |
| FGG | -1.143291 | -0.024732 | -0.29781 | 0.820751 | -0.71911 | 0.76066 | -1.07800 | -0.31734 | -0.358888 | 0.401775 |
| FN1 | -0.279561 | -0.315325 | 0.28991 | 0.254146 | -0.57776 | 1.14627 | -1.25508 | -0.10880 | -0.677313 | 0.468960 |
| GC | -0.558347 | 0.405063 | -0.79501 | 0.168400 | -1.87002 | -0.29614 | -1.26410 | -1.56024 | 0.605915 | 0.309780 |
| GSN | 0.070485 | 0.047944 | -0.67096 | -0.693498 | — | — | — | — | — | — |
| HABP2 | — | — | — | — | -0.53672 | 1.44460 | -0.70709 | 0.73751 | -0.170366 | 1.274230 |
| HP | -1.246860 | 0.527621 | -0.34881 | 1.425674 | -0.63935 | 0.96834 | -1.29633 | -0.32799 | -0.656978 | 0.311361 |
| HPX | -0.410464 | -0.288078 | -0.71149 | -0.589104 | -0.35977 | 0.93602 | -1.10344 | -0.16741 | -0.743669 | 0.192356 |
| HRG | 0.597903 | 1.067289 | 0.03216 | 0.501543 | -0.73007 | 0.68937 | -0.82317 | -0.13380 | -0.093096 | 0.596274 |
| IGHA1 | 1.763588 | 1.347662 | 0.36289 | -0.053035 | -2.01524 | 0.43280 | -2.20811 | -1.77531 | -0.192874 | 0.239928 |
| IGHD | — | — | — | — | -2.44996 | 0.41823 | -3.42847 | -3.01025 | -0.978509 | -0.560281 |
| IGHG1 | -0.085531 | 0.929213 | -0.49630 | 0.518448 | -0.09702 | -1.80911 | 0.48143 | -1.32767 | 0.578457 | -1.230650 |
| IGHG2 | 0.972042 | 0.350168 | 0.46080 | -0.161075 | -0.62494 | -1.51067 | 0.27055 | -1.24013 | 0.895491 | -0.615183 |
| IGHG3 | -0.194151 | 1.432323 | -0.93099 | 0.695486 | -1.85435 | -0.39273 | -1.88702 | -2.27975 | -0.032671 | -0.425399 |
| IGHM | -0.631813 | -0.896730 | -0.41747 | -0.682387 | -1.17417 | 1.79160 | -2.35087 | -0.55927 | -1.176697 | 0.614902 |
| IGKC | -0.069746 | 0.042036 | -0.11503 | -0.003249 | -1.18684 | -0.28755 | -1.17653 | -1.46407 | 0.010319 | -0.277230 |
| IGKV3D-20 | — | — | — | — | -0.36993 | -0.05368 | 0.21148 | 0.15780 | 0.581410 | 0.527728 |
| ITIH1 | -0.976657 | 0.705713 | -0.52118 | 1.161195 | -0.61492 | 0.54957 | -0.50394 | 0.04563 | 0.110982 | 0.660550 |
| ITIH2 | -0.314269 | -0.528321 | -0.23633 | -0.450384 | -0.74315 | 0.67572 | -1.21366 | -0.53794 | -0.470504 | 0.205218 |
| ITIH3 | -0.545603 | 0.613890 | 0.35127 | 1.510762 | -2.05644 | 1.29023 | -1.87432 | -0.58408 | 0.182118 | 1.472352 |
| ITIH4 | -0.066954 | -0.218936 | 0.38087 | 0.228885 | -1.08437 | 0.97731 | -1.81985 | -0.84254 | -0.735475 | 0.241832 |
| KLKB1 | — | -2.209308 | — | -0.271360 | — | — | — | — | — | — |
| KNG1 | -0.619816 | -0.002533 | -0.06763 | 0.549656 | -0.66441 | 0.80529 | 0.03123 | 0.83652 | 0.695635 | 1.500923 |
| LRG1 | -0.798801 | 0.256510 | 0.14022 | 1.195530 | -0.95160 | 1.70177 | -2.19510 | -0.49334 | -1.243508 | 0.458260 |
| LUM | 0.083232 | 0.658010 | -1.26356 | -0.688779 | — | — | — | — | — | — |
| ORM1 | -0.197477 | 1.117819 | -0.22401 | 1.091281 | -1.91264 | 1.67614 | -1.30260 | 0.37354 | 0.610042 | 2.286181 |
| PGLYRP2 | — | — | — | — | — | — | — | — | — | — |
| PLG | -0.368007 | 0.088056 | -0.84104 | -0.384974 | -1.07016 | 2.71125 | -2.84933 | -0.13808 | -1.779167 | 0.932079 |
| PROS1 | -0.330086 | 0.062396 | -0.79634 | -0.403862 | -0.50896 | 1.53496 | -3.87453 | -2.33957 | -3.365566 | -1.830603 |
| RBP4 | 0.450569 | 0.418579 | -0.02117 | -0.053164 | -4.09712 | 1.43523 | -2.98773 | -1.55250 | 1.109395 | 2.544623 |
| SAA1 | -2.777812 | 2.346357 | -0.51519 | 4.608983 | -1.38588 | 2.48550 | -2.55939 | -0.07388 | -1.173506 | 1.311999 |
| SERPINA1 | 0.682559 | 0.048200 | 1.78242 | 1.148065 | -0.09991 | -0.15590 | -1.36351 | -1.51941 | -1.263595 | -1.419492 |
| SERPINA3 | -0.758237 | -0.161767 | 0.18370 | 0.780166 | -0.74175 | 2.23111 | -2.03535 | 0.19576 | -1.293593 | 0.937517 |
| SERPINA4 | 0.009912 | — | -1.01801 | — | -1.44737 | — | -0.65715 | — | 0.790218 | — |
| SERPINA5 | — | — | — | 0.275703 | — | — | — | — | — | — |
| SERPINC1 | -0.555349 | -0.233936 | -0.54212 | -0.220711 | -0.77203 | 1.10667 | -1.34645 | -0.23978 | -0.574424 | 0.532243 |
| SERPIND1 | 0.253612 | — | 0.04593 | — | 0.30501 | 2.38443 | -1.64689 | 0.73754 | -1.951891 | 0.432539 |
| SERPING1 | -1.161476 | 0.119157 | -1.35109 | -0.070457 | -0.93019 | 1.07668 | -1.09046 | -0.01378 | -0.160275 | 0.916406 |
| TF | -0.282363 | -0.110509 | -0.48437 | -0.312514 | -0.76819 | 0.58757 | -0.99456 | -0.40699 | -0.226372 | 0.361200 |
| VTN | -0.618610 | -0.032377 | -0.26900 | 0.317232 | -1.72346 | 1.49185 | -2.15176 | -0.65991 | -0.428298 | 1.063555 |
| VWF | — | 1.058575 | — | 1.391788 | -2.56629 | 0.51616 | -1.97740 | -1.46124 | 0.588889 | 1.105052 |

**Table S2:**
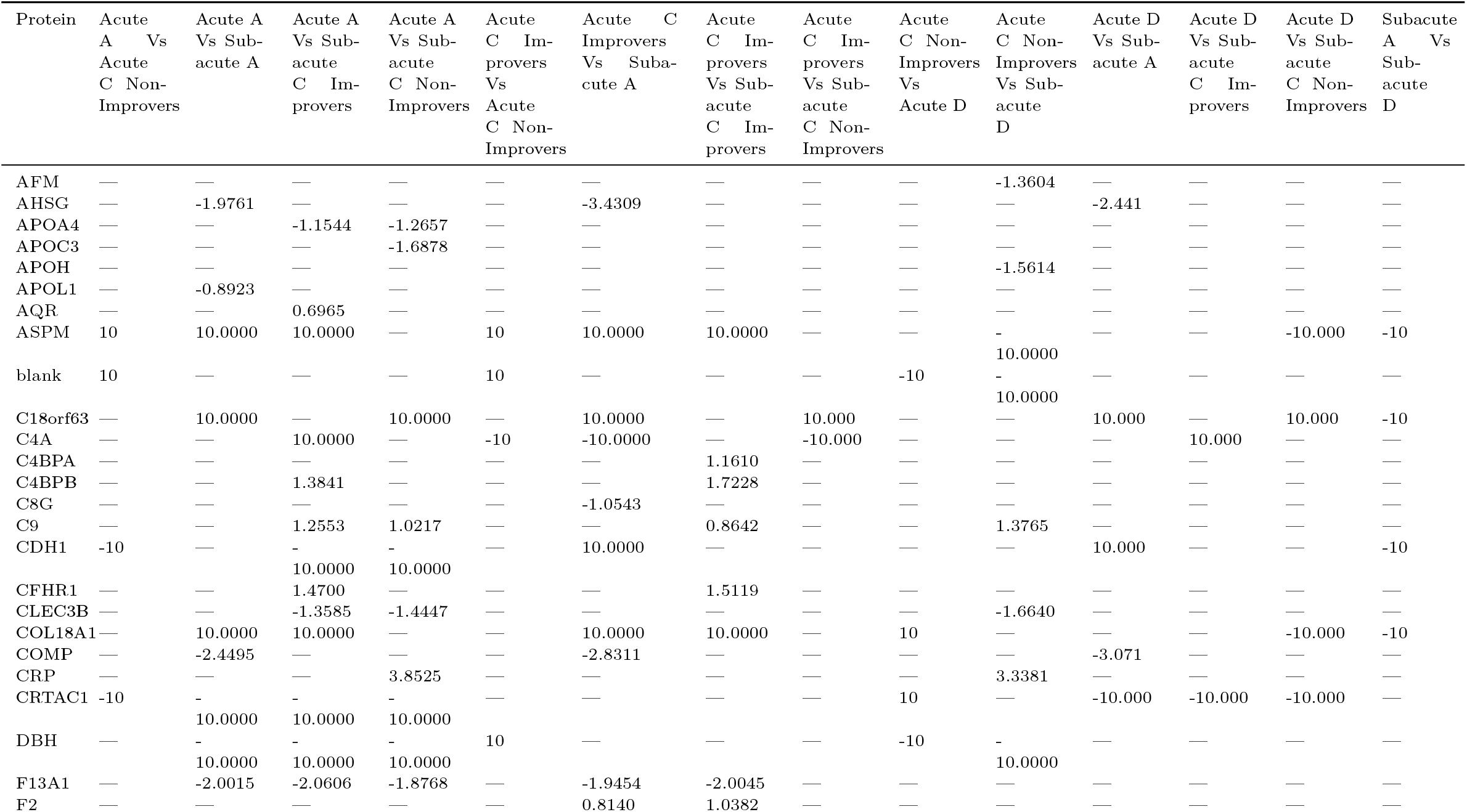
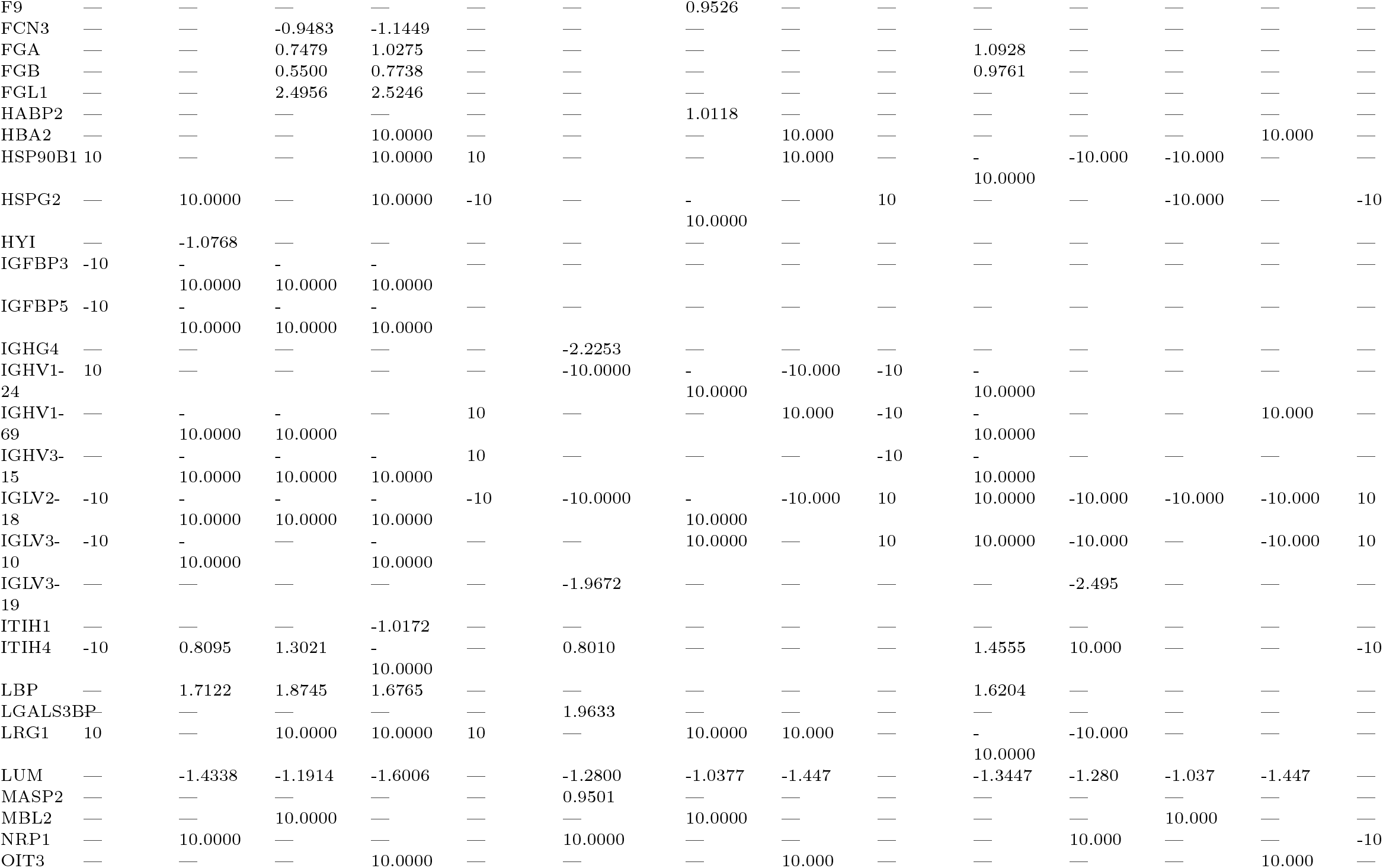
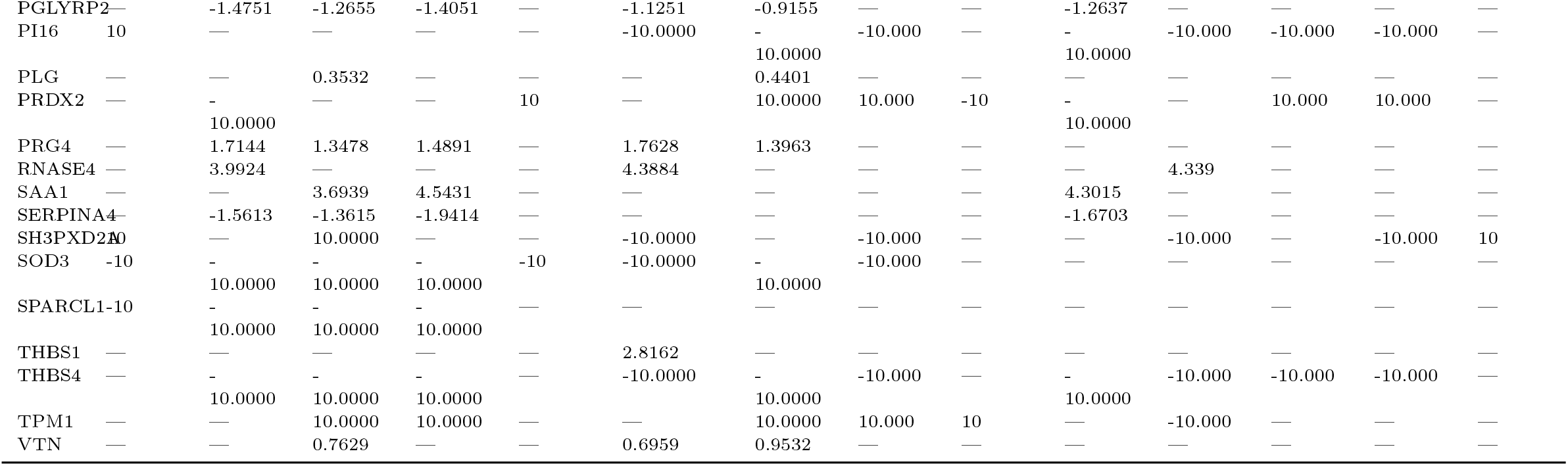
OpenMS log_2_ fold changes in the plasma proteome of SCI patients from label-free experiments. ‘Acute’ and ‘Subacute’ samples collected within 2 week and approximately 3-months post-injury respectively.

**Table S3:**
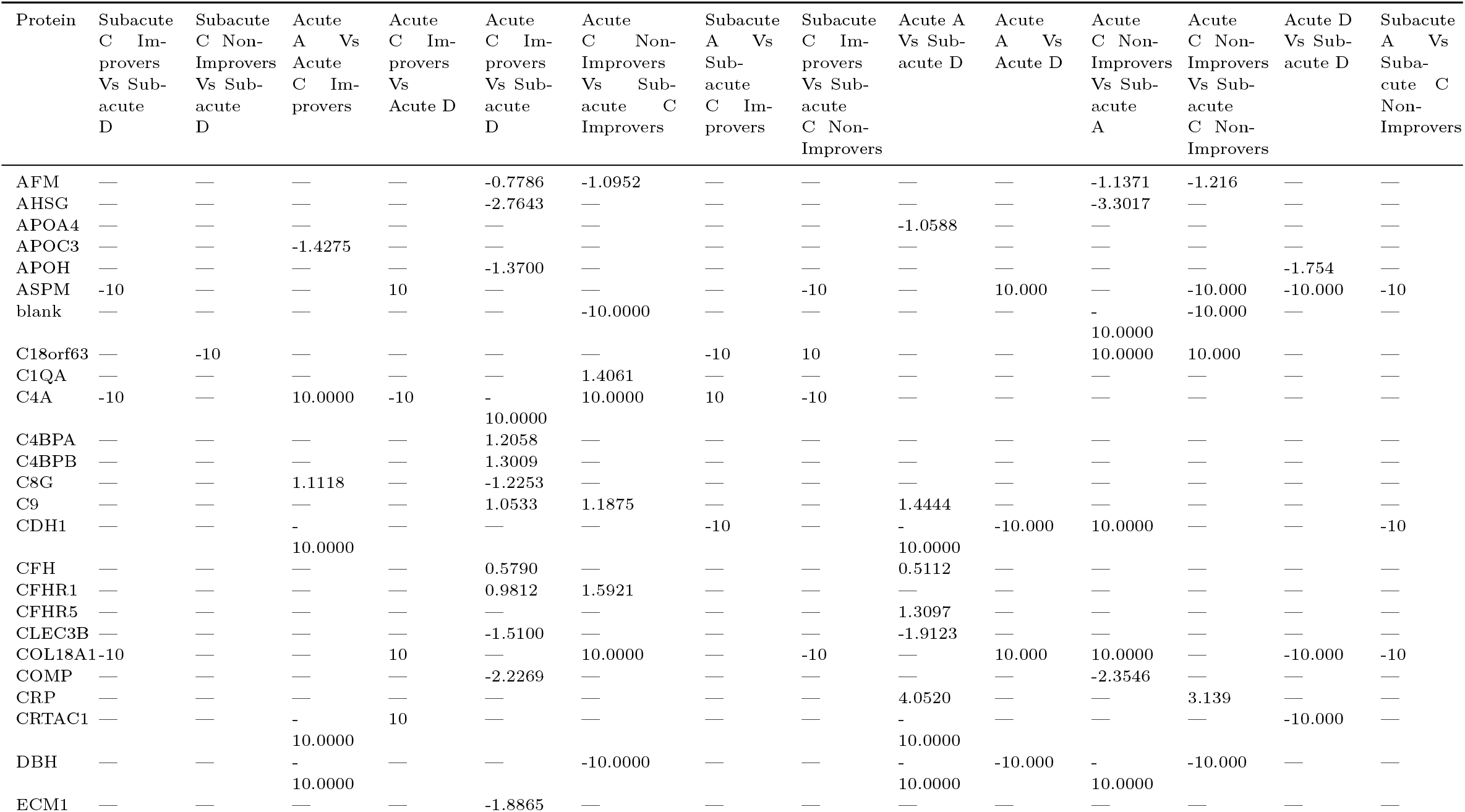
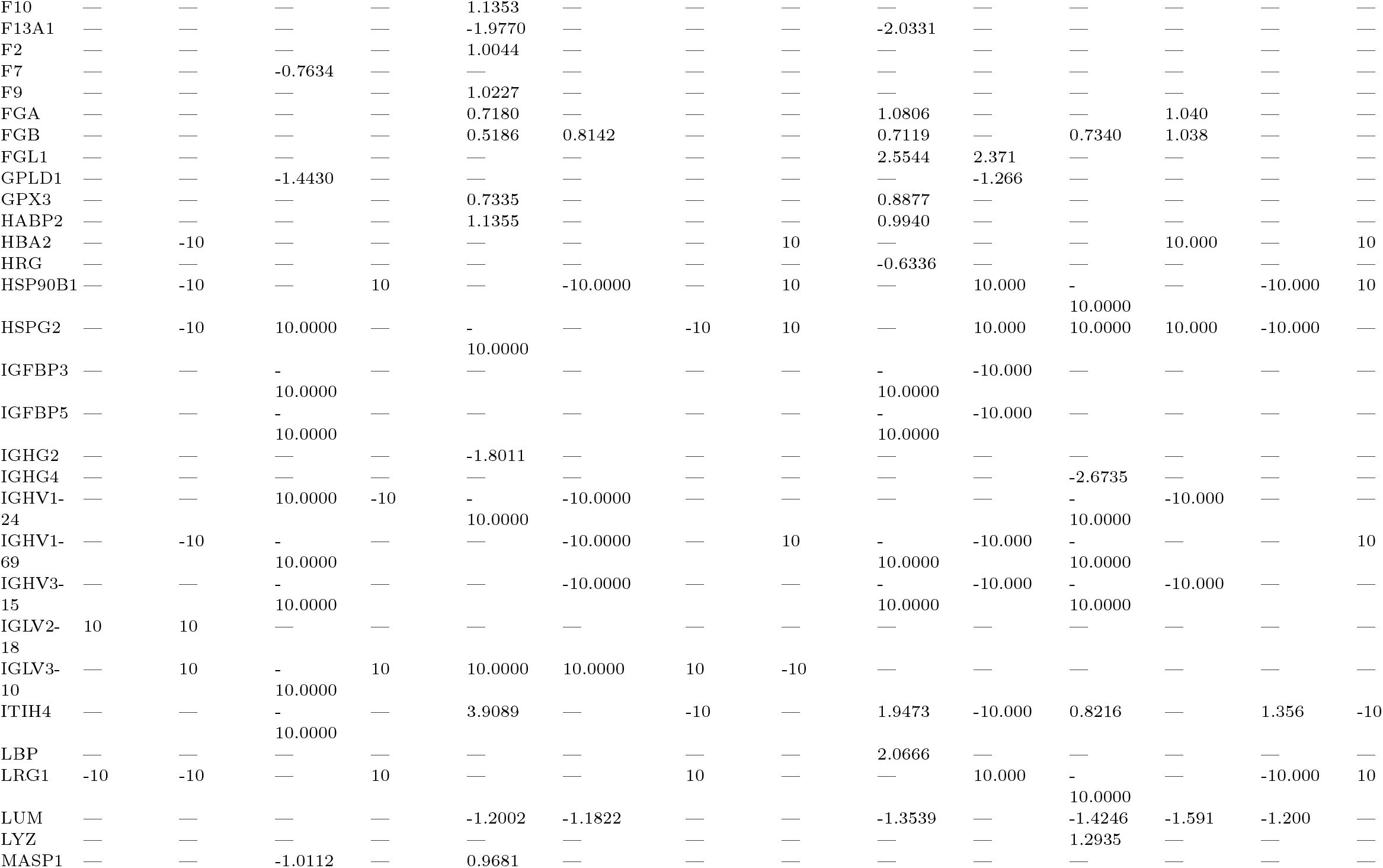
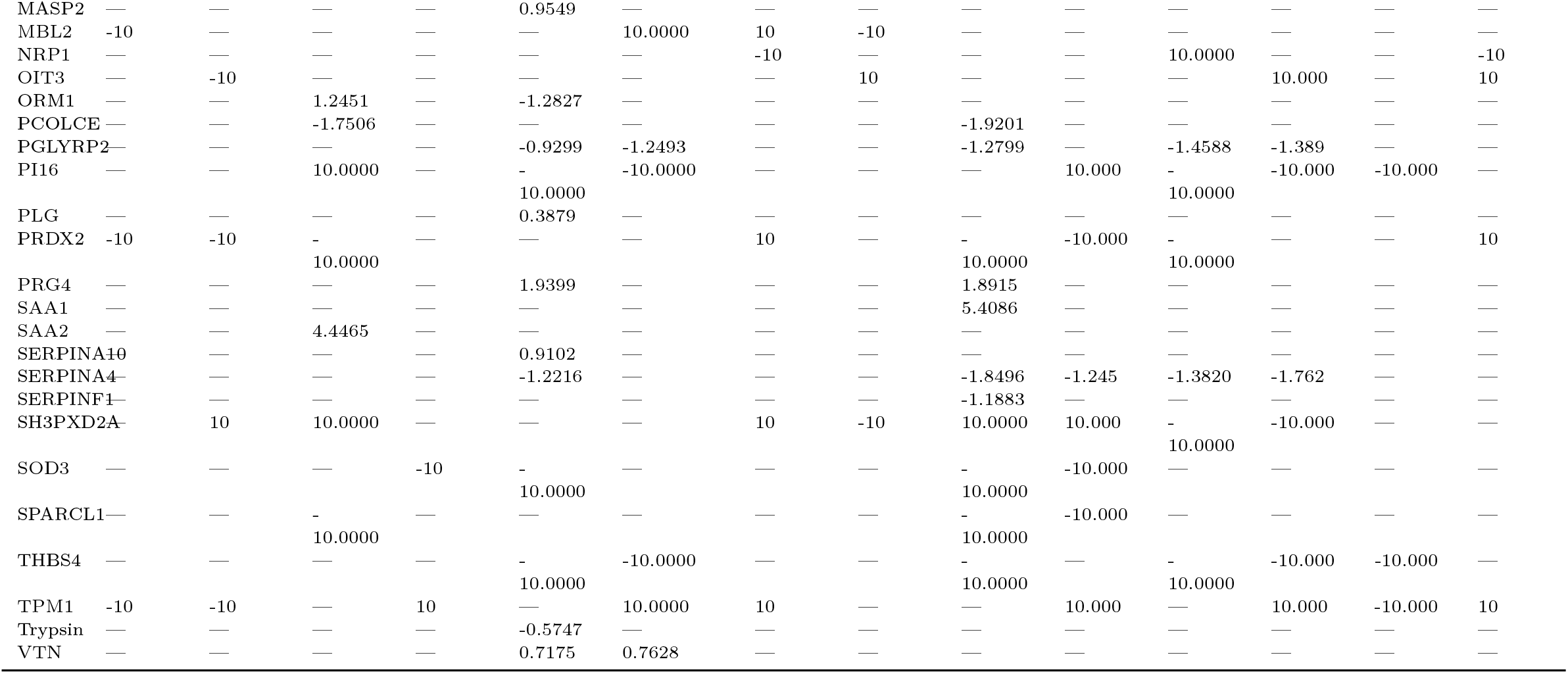
OpenMS log_2_ fold changes in the plasma proteome of SCI patients from label-free experiments. ‘Acute’ and ‘Subacute’ samples collected within 2 week and approximately 3-months post-injury respectively.

## Notes

### Competing Interest Statement

The authors have declared no competing interest.

### Summary of Updates

Coding error resulting in some proteins not being included in supplementary tables. This has been fixed, in addition to some typos.

